# Calcium signals tune AMPK activity and mitochondrial homeostasis in dendrites of developing neurons

**DOI:** 10.1101/2022.12.27.521995

**Authors:** Akane Hatsuda, Junko Kurisu, Kazuto Fujishima, Ayano Kawaguchi, Nobuhiko Ohno, Mineko Kengaku

## Abstract

The growth of dendritic arbors in developing neurons is known to be regulated by neuronal activity, which is thought as one of the mechanisms of neural circuit optimization. Here we demonstrate that activity-dependent calcium signaling controls mitochondrial homeostasis and dendritic outgrowth via AMP-activated protein kinase (AMPK) in developing hippocampal neurons. AMPK deficiency phenocopies the inhibition of neuronal activity, inducing dendritic hypotrophy with abnormally elongated mitochondria. In growing dendrites, AMPK is activated by neuronal activity and dynamically oscillates in synchrony with calcium spikes, and this AMPK oscillation is inhibited by CaMKK2 knockdown. AMPK activation leads to phosphorylation of MFF and ULK1, which initiate mitochondrial fission and mitophagy, respectively. Dendritic mitochondria in AMPK-depleted neurons exhibit impaired fission and mitophagy and display multiple signs of dysfunction. Thus, AMPK activity is finely tuned by the calcium-CaMKK2 pathway and regulates mitochondrial homeostasis by facilitating removal of damaged components in rapidly growing neurons during normal brain development.

## Introduction

Dendrite morphogenesis is a pivotal step for the establishment of neural circuit connectivity in the developing brain. It is widely accepted that Ca^2+^ signals evoked by neuronal activity play important roles in dendrite development. Ca^2+^ influx via voltage-gated calcium channels (VGCCs) and NMDA receptors triggers activation of Ca^2+^/calmodulin-dependent kinases (CaMKs) that positively and negatively regulate dendrite growth through transcriptional control and local remodeling of the cytoskeleton^1–3^.

Rapid extension and branching of dendrites in developing neurons is accompanied by extensive cytoskeletal and membrane remodeling and intracellular transport that increase the energetic burden particularly in growing dendritic branches. We have previously shown that mitochondria are actively transported to growing dendritic arbors to maintain local ATP levels necessary for the regulation of actin dynamics^4^. In addition to producing the majority of cellular ATP, mitochondria also participate in various processes indispensable for the development and function of dendrites, including Ca^2+^ homeostasis, lipid flux, and reactive oxygen species (ROS) generation^5,6^. The abnormal morphology of dendritic mitochondria is one of the pathogenic hallmarks of Alzheimer’s disease (AD), supporting the notion that constant delivery of functional mitochondria throughout a large volume of dendrites is critical for maintenance as well as differentiation of neurons^7,8^.

Mitochondrial homeostasis in postmitotic neurons is highly dependent on dynamic fission and fusion events. Mitochondrial fission is particularly important for the biogenesis of new mitochondria and for the removal of damaged or aged mitochondria through mitophagy^9^. Mitochondrial fission is triggered by the spiral assembly of an evolutionally conserved GTPase dynamin-related protein 1 (Drp1) around the constriction site on the mitochondrial surface. During fission, Drp1 is recruited to the outer mitochondrial membrane by one of its receptors, such as Mitochondrial Fission Factor (MFF)^10–12^. Several groups, including us, have shown that the inhibition of mitochondrial fission leads to abnormal elongation and dysfunction of mitochondria, resulting in aberrant dendrite formation in vivo and in vitro^13–15^. On the other hand, excessive mitochondrial fission also leads to reduced respiratory activity and abnormal distribution of mitochondria, which impairs normal development of dendrites^16,17^. However, it remains unclear how the proper balance of mitochondrial fission is adaptively regulated in developing neurons whose shape, volume and activity states are dynamically changing.

The balance between fission and fusion should be finely tuned in response to changes in the cellular and subcellular metabolic state. Recent studies have highlighted the role of AMP-activated protein kinase (AMPK) as an energy sensor that sustains the metabolic state by regulating mitochondrial dynamics and functions^18^. AMPK regulates several aspects of mitochondrial homeostasis by direct phosphorylation of key factors, including MFF, peroxisome proliferator-activated receptor-coactivator 1 (PGC1α), Armadillo repeat-containing protein 10 (ARMC10), Unc-51 like autophagy activating kinases 1 and 2 (ULK1/2) and Mitochondrial fission regulator 1-like protein (MTFR1L)^19–25^. AMPK is activated by direct allosteric binding of AMP, facilitating phosphorylation at Thr172 by an upstream kinase LKB1 in response to cellular ATP shortage^26^. Additionally, AMPK can be directly phosphorylated by the calcium (Ca^2+^)- sensitive kinase CaMKK2 (also known as CaMKKβ) in response to an elevated intracellular Ca^2+^ concentration^27,28^. In the central nervous system, AMPK has been involved in both the prevention and progression of neurodegenerative diseases, including AD. AMPK is activated by metabolic and oxidative stresses in early pathogenesis of AD, improving mitochondrial functions to restore energy balance^29^. On the other hand, overactivation of the CaMKK2-AMPK pathway by excessive Ca^2+^ signals has been implicated in mitochondrial fragmentation and degradation during the pathogenesis of Alzheimer’s disease^30,31^. However, AMPK function is less understood in the differentiating neurons during normal brain development.

In this study, we investigated the impact of neuronal activity on the mitochondrial dynamics in differentiating hippocampal neurons. We demonstrate that neuronal activity finely tunes the CaMKK2-AMPK pathway in growing dendrites. Normal AMPK activity induces mitochondrial fission and mitophagy via phosphorylation of MFF and ULK1, which are critical to remove dysfunctional mitochondrial components and maintain functional mitochondria in growing dendrites. Thus, there is indeed a causal link between activity-dependent dendritic arbor development and AMPK-dependent mitochondrial quality control.

## Results

### Neuronal activity enhances dendrite formation and mitochondrial fission in developing hippocampal neurons

We previously established long-term live-imaging of dissociated hippocampal neurons and observed the gradual formation of dendritic arbors, with dynamic extension and retraction, until the second week of culture^32^. To visualize neuronal activity in developing dendrites, we transfected GCaMP6s in cultured hippocampal neurons and performed live imaging at 5 days in vitro (DIV5). We observed intermittent Ca^2+^ transients in dendrites and cell soma, which were abolished by the combined treatment with a blocker of voltage-gated sodium channels, tetrodotoxin, and an NMDA receptor antagonist, D-(-)-2-Amino-5-phosphonopentanoic acid (TTX+APV treatment) (Fig. 1A, Supplementary Video 1). There was no apparent difference in the dynamic extension and retraction of dendrites in drug-treated and untreated neurons, but growth retardation gradually became evident, with a significant decrease in total dendritic length and the number of branches at 48 hr after TTX+APV treatment (Fig. 1B-D, Supplementary Video 2).

**Figure 1.**
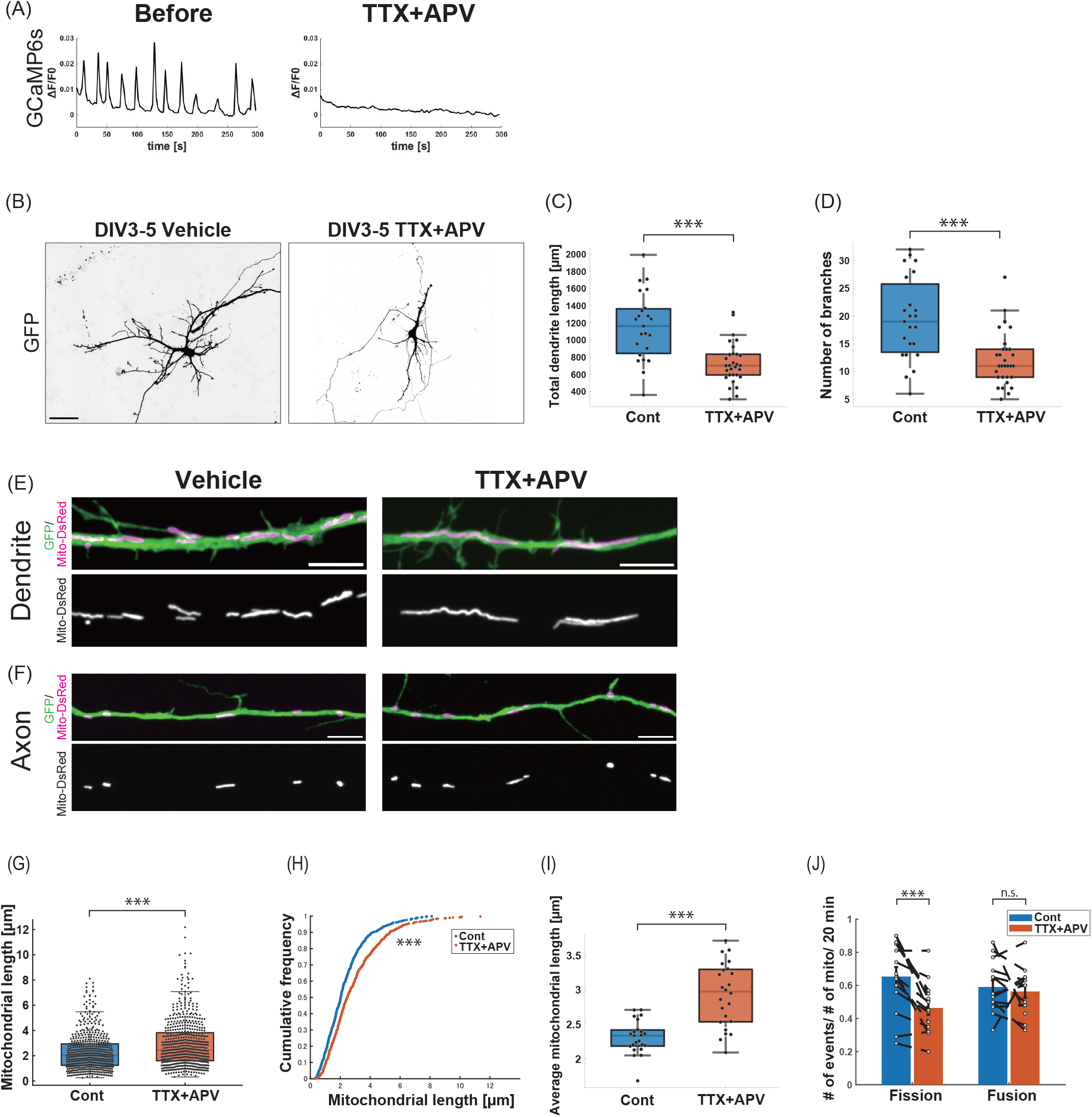
Neuronal activity enhances dendrite formation and mitochondrial fission in developing hippocampal neurons. **(A)** GCAMP6s ΔF/F transients in cultured hippocampal neurons before and after TTX+APV treatment (see also Supplementary Video 1). **(B)** Morphology of hippocampal neurons treated with vehicle or TTX+APV from DIV3 to 5. Cells were labeled with GFP (see also Supplementary Video 2). **(C, D)** Quantification of total dendritic length (C) and number of branch points (D). N= 23 for control cells, 30 for TTX+APV treated cells. ***p= 4.8×10^−5^ (C); ***p= 1.8×10^−4^ (D); Wilcoxon rank-sum test. **(E, F)** Morphology of mitochondria in dendrites (E) and axons (F) of DIV5 hippocampal neurons treated with vehicle or TTX+APV for 6h. Cells were labeled with mito-DsRed and GFP. **(G, H)** Distribution (G) and cumulative frequency (H) of the mitochondrial length in dendrites. N= 576 mitochondria from 23 control cells, 613 mitochondria from 24 drug-treated cells. ***p= 1.3×10^−9^, Wilcoxon rank-sum test. **(I)** Average length of dendritic mitochondria in individual neurons. N= 23 control cells, 24 TTX+APV treated cells. ***p= 1.2×10^−5^, Wilcoxon rank-sum test. **(J)** Quantification of the mitochondrial fission and fusion frequencies in dendrites before and 1-3h after TTX+APV treatment. N= 14 cells, mean± SEM, ***p= 4.1×10^−4^ for fission, (n.s.) p= 0.57 for fusion, paired two-tailed t test. Box plots denote the median (50th percentile) and the 25th to 75th percentiles of datasets in (C, D, G, I). Samples were collected from three independent experiments. Scale bars: 50 μm in (B), 5 μm in (E, F).

To examine if neuronal activity affects mitochondrial dynamics in growing dendrites, we transfected Mito-DsRed into neurons that were then untreated or treated with TTX+APV. Under both conditions, mitochondria were distributed throughout the dendritic arbors, and a significant fraction of them moved along the dendritic shaft as previously reported by us and others^4,33–35^. Inhibition of neuronal activity did not affect mitochondrial motility in dendrites: neither the fraction of motile mitochondria nor their average speed was altered by TTX+APV treatment (Extended Data Fig. 1A and B). There were no apparent changes in the density and distribution of mitochondria in dendrites in the drug-treated neurons (Extended Data Fig. 1C). However, we found a significant increase in the length of dendritic mitochondria in the drug-treated neurons [Median (IQR)= 2.0 (1.2-2.9) μm in control vs 2.5 (1.6-3.8) μm in TTX+APV neurons; Fig. 1E, G, H, I]. In the axon, on the other hand, the length of mitochondria was much shorter, as previously reported^36^, and was unaffected by TTX+APV treatment (Fig. 1F). As mitochondrial morphology is affected by the balance between fission and fusion, we analyzed the frequency of fission and fusion in dendrites by high resolution time-lapse imaging (Extended Data Fig. 1D and Supplementary Video 3). We found that inhibition of neuronal activity significantly suppressed mitochondrial fission but had little or no effect on fusion (Fig. 1J). Conversely, glutamate treatment led to a marked increase in smaller mitochondria in the cell soma and dendrites, suggesting that enhanced neuronal activity facilitated mitochondrial fission (Extended Data Fig. 1E). Taken together, these data suggest that neuronal activity is involved in the regulation of the mitochondrial fission-fusion balance during dendritic outgrowth in developing neurons.

### AMPK regulates dendritic growth and mitochondrial fission

It has been shown that AMPK mediates mitochondrial fission in response to energy stress in neurons and non-neuronal cells^20^. We therefore examined if AMPK is involved in activity-dependent mitochondrial fission in developing hippocampal neurons. AMPK is a heterotrimeric serine/threonine kinase composed of one catalytic (α1/α2), one regulatory (β1/β2) and one AMP/ATP binding (γ1/γ2/γ3) subunit. Both AMPKα1 and AMPKα2 were expressed in cultured hippocampal neurons, as assessed by western blotting using isoform-specific antibodies (Fig. 2A). Specifically, AMPKα1 expression was high in earlier stages and declined by DIV10, whereas AMPKα2 expression started at a level comparable to AMPKα1, and then gradually increased, peaking at DIV10. The abundance of active AMPKα phosphorylated at Thr172 (detected by an antibody that recognizes both phosphorylated isoforms, p-AMPKα) increased during dendritic outgrowth in parallel with AMPKα2 expression (Fig. 2A).

**Figure 2.**
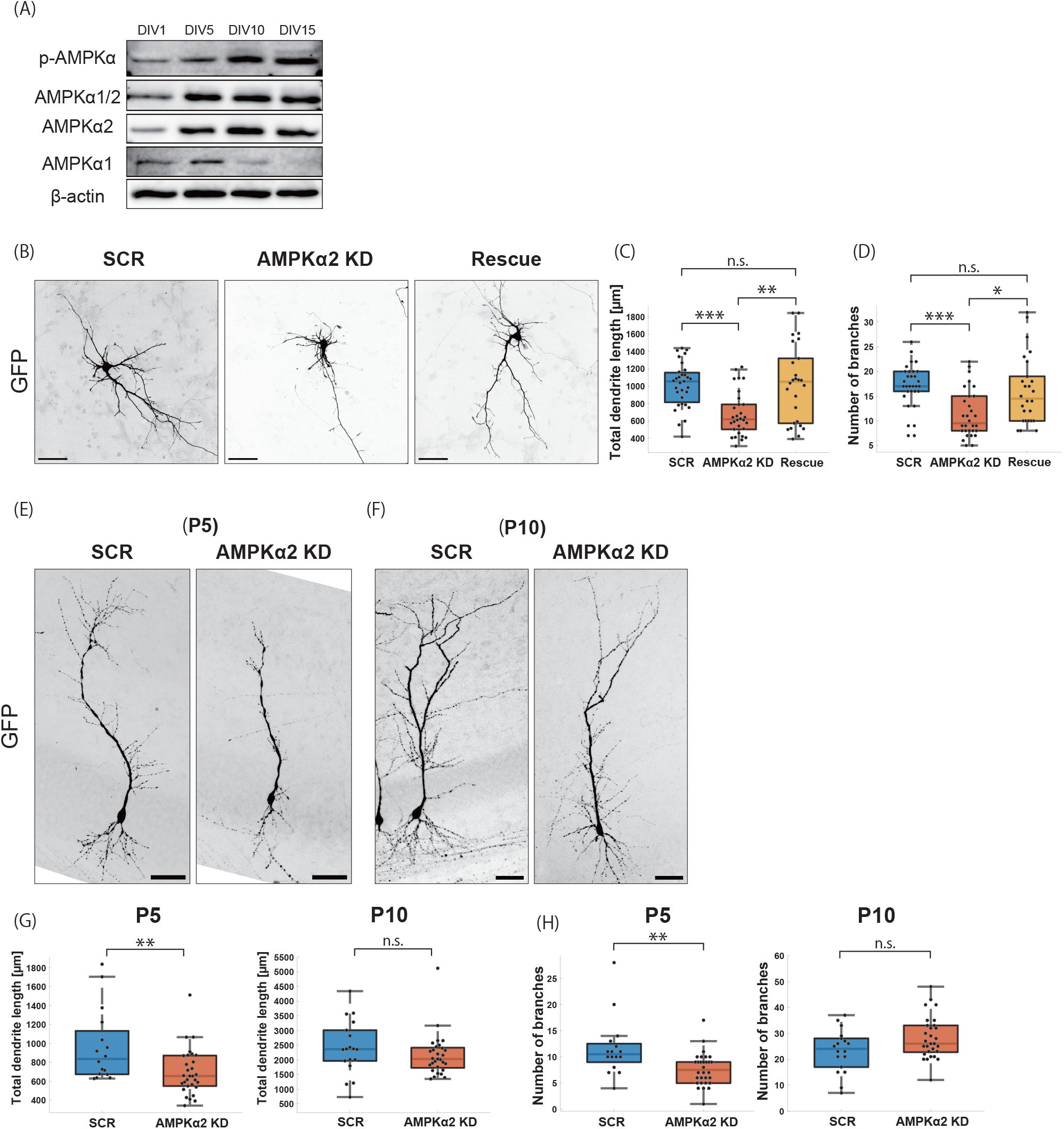
AMPK promotes dendrite formation of hippocampal neurons. **(A)** Expression profiles of AMPKα subunits in developing hippocampal neurons in culture. Whole cell lysates were collected at indicated stages and subjected to western blotting with antibodies against AMPKα1 and/or α2, and AMPKα1/2 phosphorylated at threonine 172 (p-AMPKα). **(B)** Morphology of hippocampal neurons transfected with GFP and the following constructs: shRNA-scramble control (SCR); shRNA-AMPKα2 #1 (AMPKα2 KD); or shRNA-AMPKα2 #1 with shRNA-resistant AMPKα2 (Rescue). Cells were transfected at DIV3 and fixed at DIV5. **(C, D)** Quantification of total dendritic length (C) and number of branch points (D). N= 31 cells for SCR, 30 cells for AMPKα2 KD, and 26 cells for Rescue. ***p= 4.0×10^−4^ for SCR vs. AMPKα2 KD, **p= 0.0058 for AMPKα2 KD vs. Rescue (C); ***p= 2.0×10^−4^ for SCR vs. AMPKα2 KD, *p= 0.018 for AMPKα2 KD vs. Rescue (D), Kruskal–Wallis test with Bonferroni multiple comparisons test. **(E, F)** Morphology of CA1 pyramidal neurons transfected with GFP and shRNA-scramble control (SCR) or shRNA-AMPKα2 #1 (AMPKα2 KD) by in utero electroporation at E14.5, followed by fixation at P5 (E) and P10 (F). **(G, H)** Quantification of total dendritic length (G) and number of branch points (H) in CA1 pyramidal neurons at P5 and P10. N≥ 16 cells for SCR and AMPKα2 KD, **p= 0.0067 for P5; (n.s.) p= 0.20 for P10 (G); **p= 0.0021 for P5; (n.s.) p= 0.12 for P10 (H), Wilcoxon rank-sum test. Box plots denote the median (50th percentile) and the 25th to 75th percentiles of datasets in (C, D, G, H). Samples were collected from three independent experiments. Scale bars: 50 μm in (B, E, F).

To investigate the role of AMPK in mitochondrial dynamics and dendritic formation, we knocked down AMPK subunits in cultured neurons by expressing short hairpin RNAs (shRNA) against AMPKα1 and AMPKα2 (Extended Data Fig. 2A and B). At DIV3, we transfected dissociated hippocampal neurons with AMPKα1 shRNA or AMPKα2 shRNA plasmids together with a volume marker GFP construct, and observed dendritic morphology at DIV5. AMPKα2 shRNA significantly reduced the size and complexity of dendritic branches, reminiscent of the activity-deprived neurons (Fig. 2B-D and Extended Data Fig. 2C-E compared to Fig. 1). Concomitant exogenous expression of an AMPKα2 mutant cDNA (AMPKα2 Rescue) that was resistant to AMPKα2 shRNA largely prevented the dendritic defects, discounting any major off-target effects of the shRNA (Fig. 2B-D and Extended Data Fig. 2C). In contrast, AMPKα1 shRNA produced much weaker effects on dendritic morphology. Further, simultaneous knockdown of AMPKα1 and α2 was not phenotypically different from AMPKα2 knockdown, suggesting that AMPKα2 was the predominant isoform regulating dendritic morphogenesis in our culture (Extended Data Fig. 2C-E). We further confirmed that repression of AMPKα2 using CRISPR interference (CRISPRi) significantly delayed dendritic outgrowth at DIV5 (Extended Data Fig. 2F-H). We thus focused on AMPKα2 in subsequent experiments.

To confirm the requirement of AMPKα2 for dendritic formation in vivo, we introduced the AMPKα2 shRNA#1 together with GFP into CA1 pyramidal neurons at embryonic day (E) 14.5 by in utero electroporation. Compared to control neurons, AMPKα2-depleted neurons showed significantly reduced length and complexity of the apical dendrite at P5, supporting the notion that AMPKα2 functions in dendritic outgrowth of hippocampal pyramidal neurons (Fig. 2E-H). In contrast, the dendritic hypotrophy was less evident at P10, suggesting a compensatory mechanism allows the recovery of dendritic growth retardation in the context of AMPKα2 deficiency in vivo.

Returning to cultured cells, we next observed mitochondria in AMPKα2-depleted neurons by cotransfecting mito-DsRed and GFP. The AMPKα2 knockdown significantly increased mitochondrial length in dendrites [Median (IQR)= 2.1 (1.3-3.1) μm in control; 2.6 (1.6-4.1) μm in AMPKα2 knockdown; 2.0 (1.3-3.1) μm in rescue; Fig. 3A, C-E], but not in the axon at DIV5 (Fig. 3B). There was no apparent change in the density and distribution of mitochondria in the hypotrophic dendrites in AMPKα2-depleted neurons (Extended Data Fig. 3A). AMPKα2 CRISPRi also induced abnormal elongation of mitochondria in dendrites (Extended Data Fig. 3B-E). These results prompted us to use time-lapse imaging to analyze the fission/fusion frequency of dendritic mitochondria in the neurons transfected with AMPKα2 shRNA. As in the case for neuronal activity inhibition, AMPKα2 knockdown specifically lowered the frequency of fission but had little effect on fusion in dendrites (Fig. 3F).

**Figure 3.**
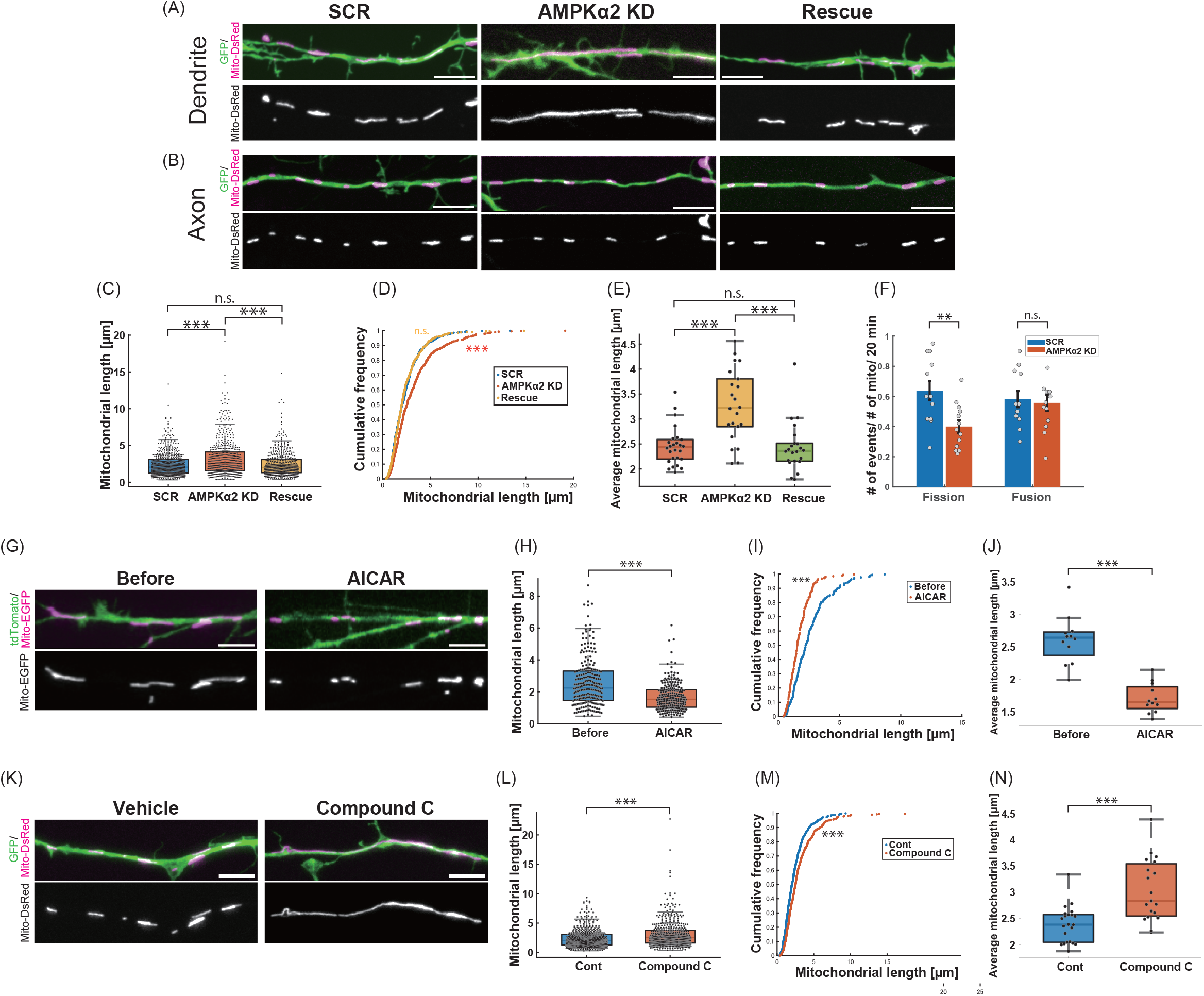
AMPK enhances mitochondrial fission in dendrites of developing hippocampal neurons. **(A, B)** Morphology of mitochondria in dendrites (A) and axons (B) in hippocampal neurons transfected with GFP and mito-DsRed together with shRNA-scramble control (SCR), shRNA-AMPKα2#1 (AMPKα2 KD), or shRNA-AMPKα2#1 with shRNA-resistant AMPKα2 (Rescue). Cells were transfected at DIV3 and fixed at DIV5. **(C, D)** Distribution (C) and cumulative frequency (D) of the length of dendritic mitochondria in SCR, AMPKα2 KD and Rescue cells. N= 669 mitochondria from 26 cells (SCR), 548 mitochondria from 22 cells (AMPKα2 KD), and 543 mitochondria from 22 cells (Rescue), respectively. ***p< 0.001; (n.s.) p> 0.05; Kruskal–Wallis test with Bonferroni multiple comparisons test. **(E)** Average length of dendritic mitochondria in individual cells. N=26 cells (SCR), 22 cells (AMPKα2 KD), and 22 cells (Rescue), respectively. ***p< 0.001; (n.s.) p> 0.05; Kruskal–Wallis test with Bonferroni multiple comparisons test. **(F)** Quantification of the mitochondrial fission and fusion frequencies in dendrites in SCR and AMPKα2 KD cells. N= 12 cells for SCR, and 13 cells for AMPKα2 KD. Mean ± SEM, **p= 0.0042; (n.s.) p= 0.75; unpaired two-tailed t test. **(G)** Morphology of dendritic mitochondria in hippocampal neurons before and 3-5 h after treatment with 1 mM AICAR (AMPK activator). Cells were labeled with tdTomato and mito-EGFP. **(H, I)** Distribution (H) and cumulative frequency (I) of the length of dendritic mitochondria before and after AICAR treatment. N= 230 mitochondria from 12 control cells, 228 mitochondria from 12 drug-treated cells, ***p= 1.9×10^−11^; Wilcoxon rank-sum test. **(J)** Average length of dendritic mitochondria in individual cells. N= 12 cells, *** p= 4.9×10^−4^, Wilcoxon signed-rank test. **(K)** Morphology of dendritic mitochondria in hippocampal neurons treated with vehicle or 20 μM Compound C for 6h. Cells were labeled with GFP and mito-DsRed. **(L, M)** Distribution (L) and cumulative frequency (M) of the length of dendritic mitochondria treated with vehicle or Compound C. N= 575 mitochondria from 22 control cells, 483 mitochondria from 19 drug-treated cells, ***p= 8.5×10^−8^, Wilcoxon rank-sum test. **(N)** Average length of the dendritic mitochondria in individual cells. N=22 control cells, 19 drug-treated cells, *** p= 2.9×10^−4^, Wilcoxon rank-sum test. Box plots denote the median (50th percentile) and the 25th to 75th percentiles of datasets in (C, E, H, J, L, N). Samples were collected from three independent experiments. Scale bars: 5 μm in (A, B, G, K).

We further documented the importance of AMPK activity in mitochondrial dynamics by pharmacological activation and inhibition. We found that dendritic mitochondria were significantly shortened after a 3-5 h treatment with an AMPK activator, AICAR (1 mM) (Fig. 3G-J). In contrast, treatment with an AMPK inhibitor, Compound C (20 μM), elongated dendritic mitochondria (Fig. 3K-N), underscoring the involvement of AMPK in the regulation of mitochondrial fission in hippocampal neurons. Collectively, these data suggest that AMPKα2 has critical roles in mitochondrial fission and dendritic formation in hippocampal neurons, and that the two phenomena are mechanistically linked.

### Neuronal activity induces dynamic oscillation of AMPK activity via Ca/CaMKK2

The striking phenotypic similarities between the modulation of neuronal activity and AMPK activity prompted us to examine if AMPK mediates the activity-dependent control of mitochondrial dynamics in developing dendrites. In order to analyze the spatiotemporal dynamics of AMPK activity in developing hippocampal neurons, we transfected them with AMPKAR-EV, a highly sensitive fluorescence resonance energy transfer (FRET)-based biosensor for AMPK^37^. We observed high basal FRET signals in the axon, which presumably reflected the activity of AMPK-related kinases such as SAD-A/B^38–40^. Indeed, AMPKα2 shRNA specifically suppressed FRET signals in the somatodendritic compartment but not in the axon (Extended Data Fig. 4A), supporting the notion that the FRET signals in dendrites, but not in axons, reflect AMPK activity.

Consistent with previous reports, we observed a rapid increase in AMPK activity in the cell body and dendrites upon glutamate treatment^41,42^; moreover, the AMPK activity was abrogated by AMPKα2 knockdown (Extended Data Fig. 4B). Interestingly, we found an unambiguous periodic fluctuation of AMPK activity with a frequency of 0 to 4 peaks in 3 minutes, in the somatodendritic compartment of untreated neurons (Fig. 4A and C). The basal fluctuation was suppressed by cotransfection of AMPKα2 shRNA or inhibition of neuronal activity by TTX+APV treatment, suggesting that neuronal activity caused dynamic changes in AMPK activity (Fig. 4A-D). To further prove the causal relationship of neuronal activity and AMPK activation, we performed simultaneous imaging of Ca^2+^ and AMPK activity using a red fluorescent Ca^2+^ indicator, jRGECO1a, together with AMPKAR-EV. We observed periodic Ca^2+^ spikes which highly synchronized with AMPK activity fluctuation in dendrites (Fig. 4E, F and Supplementary Video 4). The temporal cross-correlation analysis revealed the highest correlation at 5 sec after Ca^2+^ influx, indicating that Ca^2+^ influx precedes AMPK activation (Fig. 4G). Inhibition of neuronal activity by TTX+APV treatment suppressed both Ca^2+^ spikes and AMPK fluctuation (Extended Data Fig. 4C). In contrast, AMPK knockdown suppressed only the AMPK fluctuation, and did not alter the Ca^2+^ transient (Extended Data Fig. 4D), indicating that AMPK fluctuation was induced by neuronal activity.

**Figure 4.**
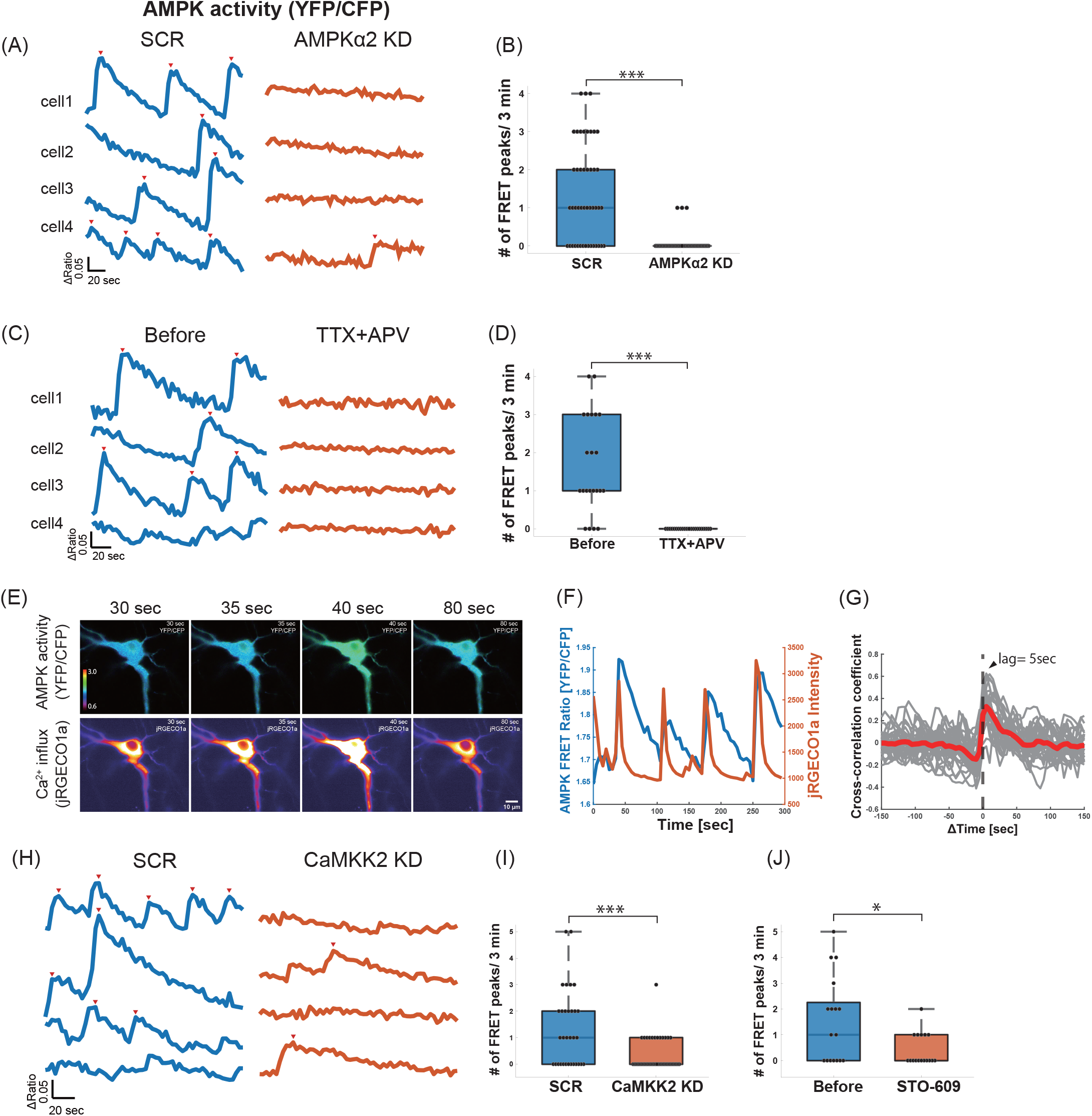
Neuronal activity induces dynamic oscillation of AMPK activity via Ca2+/CaMKK2. **(A)** Representative traces of AMPK activity (YFP/CFP ratio) in dendrites of hippocampal neurons expressing AMPKAR-EV with shRNA-scramble control (SCR) or shRNA-AMPKα2 #1 (AMPKα2 KD). Time-lapse images were acquired every 3 sec for 3 min at DIV5 using epifluorescence microscopy. Red arrowheads indicate peaks of AMPK activity. **(B)** Quantification of the frequency of AMPKAR-EV FRET peaks in hippocampal neurons described in (A). N= 46 cells for SCR, 40 cells for AMPKα2 KD; ***p= 1.2×10^−8^; Wilcoxon rank-sum test. **(C)** Representative traces of AMPK activity in hippocampal neurons expressing AMPKAR-EV before and after TTX+APV treatment. **(D)** Quantification of the frequency of AMPKAR-EV FRET peaks described in (C). N= 22 cells; ***p= 1.6×10^−4^; Wilcoxon signed-rank test. **(E, F)** Representative time-lapse images and traces of AMPKAR-EV FRET signals and Ca^2+^ influx in hippocampal neurons transfected with AMPKAR-EV and jRGECO1a. Scale bar: 10 μm (E). Blue and orange lines indicate AMPKAR-EV FRET and jRGECO1a signal intensity, respectively (F) (see also Supplementary Video 4). **(G)** The temporal cross-correlations between AMPK activity and Ca^2+^ influx. Gray lines show individual cells, and the red line indicates the average temporal cross-correlation function. The arrowhead shows the highest correlation at 5 sec after Ca^2+^ influx. N= 33 cells. **(H)** Representative traces of AMPK activity in hippocampal neurons expressing AMPKAR-EV with shRNA-scramble control (SCR) or shRNA-CaMKK2 (CaMKK2 KD). **(I)** Quantification of the frequency of AMPKAR-EV FRET peaks described in (H). N=32 cells for SCR, 45 cells for CaMKK2 KD; ***p= 6.8×10^−4^; Wilcoxon rank-sum test. **(J)** Quantification of the frequency of AMPKAR-EV FRET peaks in hippocampal neurons expressing AMPKAR-EV before and 2h after treatment with 10 μM STO-609 (CaMKK2 inhibitor). N= 17 cells from two independent experiments; *p= 0.011; Wilcoxon signed-rank test. Box plots denote the median (50th percentile) and the 25th to 75th percentiles of datasets in (B, D, I, J). Samples were taken from three independent experiments in (B, D, G, I).

Two upstream kinases, CaMKK2 and LKB1, regulate AMPK activity by phosphorylation^26–28^. As CaMKK2 has been implicated in activity-dependent formation of dendrites^43,44^, we examined whether CaMKK2 is required for AMPK fluctuation in response to neuronal activity. Cotransfection of CaMKK2 shRNA or treatment with CaMKK2 inhibitor (STO609, 10 μM) suppressed the periodic activation of AMPK (Fig. 4H-J). On the other hand, the AMPK fluctuation was not affected by treatment with an LKB1 inhibitor (Pim1, 1μM) Extended Data Fig. 4E).

Taken together, these data suggest that neuronal activity dynamically regulates AMPK activity via Ca^2+^/CaMKK2 in the somatodendritic region of developing hippocampal neurons.

### AMPKα2 knockdown prevents asymmetric mitochondrial fission and mitophagy in developing dendrites

Mitochondrial fission is involved in biogenesis and mitophagic degradation of mitochondria. It has recently been reported that the downstream pathways could be distinguished by the position of fission, either at the midzone or the periphery^45^. In normal developing hippocampal neurons, we observed both midzone and peripheral fissions in growing dendrites, with the frequency of the midzone fission, which was implicated in mitochondrial biogenesis in the previous study^45^, being approximately 2.1-fold higher than peripheral fission (Fig. 5A and B). TTX+APV treatment hampered both midzone and peripheral fissions, but had a stronger inhibitory effect on peripheral fission (Fig. 5B). AMPKα2 knockdown displayed strikingly similar effects, with stronger suppression of the peripheral fission (Fig. 5C).

**Figure 5.**
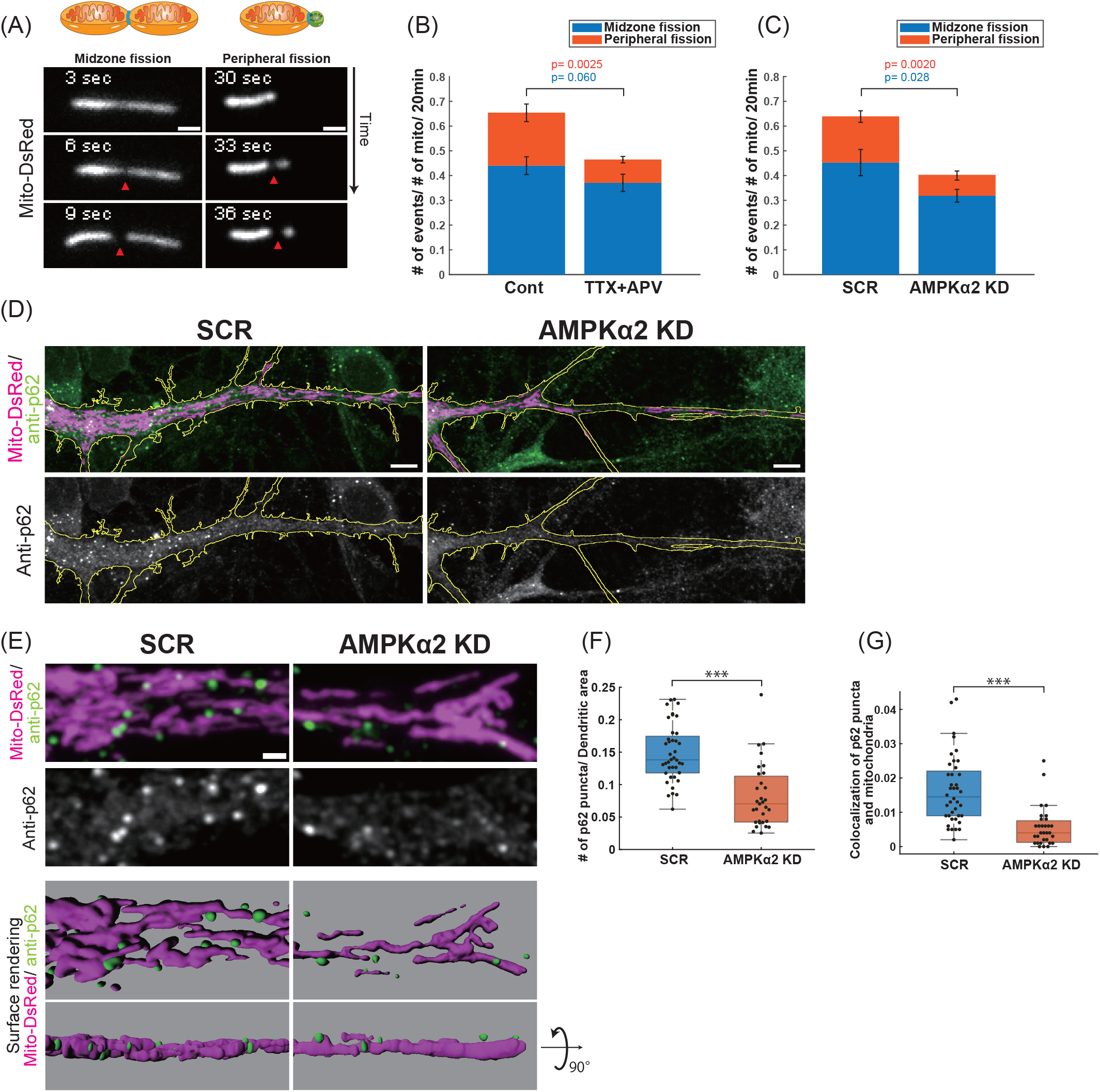
AMPK knockdown suppresses asymmetric mitochondrial fission and mitophagy in growing dendrites. **(A)** Time-lapse images of mitochondria undergoing midzone or peripheral fission in dendrites of hippocampal neurons. Cells were labelled with GFP and mito-DsRed. The red arrows indicate fission sites. **(B)** Quantification of the frequencies of midzone and peripheral fissions of dendritic mitochondria in hippocampal neurons before and after treatment with TTX+APV. N= 14 cells; mean ± SEM; (n.s.) p= 0.060 for midzone fission; **p= 0.0025 for peripheral fission; paired two-tailed t test. **(C)** Quantification of the frequencies of midzone and peripheral fissions of dendritic mitochondria in hippocampal neurons transfected with shRNA-scramble control (SCR) or shRNA-AMPKα2 #1 (AMPKα2 KD). N= 12 cells for SCR, 13 cells for AMPKα2 KD; mean ± SEM; *p= 0.028 for midzone fission; **p= 0.0020 for peripheral fission; unpaired two-tailed t test. **(D, E)** Immunofluorescence of autophagy marker p62 in dendrites of hippocampal neurons expressing shRNA-scramble control (SCR) or shRNA-AMPKα2 (AMPKα2 KD). Cells were labeled with GFP and mito-DsRed. Yellow lines indicate the outlines of dendrites (D). Higher magnified views and surface rendering images of mitochondria (magenta) and p62 puncta (green) in control and AMPKα2 KD cells (E). **(F)** Quantification of the number of p62 puncta in SCR and AMPKα2 KD cells. The number of p62 puncta was divided by the dendritic area. N= 40 cells for SCR, 31 cells for AMPKα2; ***p= 2.9×10^−7^; Wilcoxon rank-sum test. **(G)** Quantification of the co-localization (Manders overlap coefficient) of p62 puncta and mitochondrial area in SCR and AMPKα2 KD cells. N= 40 cells for SCR, 31 cells for AMPKα2; ***p= 2.1×10^−7^; Wilcoxon rank-sum test. Box plots denote the median (50th percentile) and the 25th to 75th percentiles of datasets in (F, G). Samples were taken from three independent experiments. Scale bars: 1 μm in (A, E), 5 μm in (D).

As the peripheral fission has been implicated in the mitophagy pathway, we asked whether AMPK-induced mitochondrial fission is required for mitophagic degradation of dendritic mitochondria. To this end, we used an antibody against the autophagy marker p62 to immunostain neurons transfected with Mito-DsRed with or without AMPKα2 shRNA. The p62 signals formed puncta, some of which were colocalized with mitochondria in the soma and dendrites in control neurons. By contrast, the number of p62 puncta and those colocalized with mitochondria were significantly decreased in AMPKα2 knockdown cells (Fig. 5D-G). These data suggest that AMPK-mediated mitochondrial fission is important for the mitophagy initiation during dendrite development.

We also examined the effect of AMPK activation on mitochondrial biogenesis. AMPK has been shown to phosphorylate and activate PGC1α, the master regulator of mitochondrial biogenesis. However, we detected no obvious difference in the phosphorylation level of PGC1α by Phos-tag SDS-PAGE, nor the expression of PGC1α, in response to TTX+APV, Glutamate, Compound C or AICAR (Extended Data Fig. 5A).

### AMPK phosphorylates mitochondrial fission factor and ULK1 in response to neuronal activity

AMPK phosphorylates various downstream substrates involved in many aspects of mitochondrial homeostasis. AMPK regulates mitochondrial fission by direct phosphorylation of MFF, which promotes its association with Drp1 at the mitochondrial outer membrane^19,20^. AMPK is also known to directly phosphorylate the autophagy initiating kinase ULK1^23^. We thus asked whether AMPK phosphorylates these substrates in response to neuronal activity.

Consistent with our observation that AMPK activity was dynamically regulated in synchrony with neuronal activity, AMPK phosphorylation rapidly increased within 5 min of glutamate treatment (Fig. 6A). In contrast, TTX+APV treatment reduced the basal level of AMPK phosphorylation. Concomitantly, MFF phosphorylation at Ser146 was increased by glutamate treatment, while it was repressed by TTX+APV treatment. Likewise, the level of ULK1 phosphorylation at Ser555 paralleled that of AMPK phosphorylation, which was altered by manipulation of the activity states in cultured neurons. We also confirmed that the phosphorylation levels of MFF and ULK1 were upregulated by AICAR treatment, while they were downregulated by Compound C treatment. Taken together, these results suggest that AMPK activation by neuronal activity induces mitochondrial fission and mitophagy by direct phosphorylation of MFF and ULK1.

**Figure 6.**
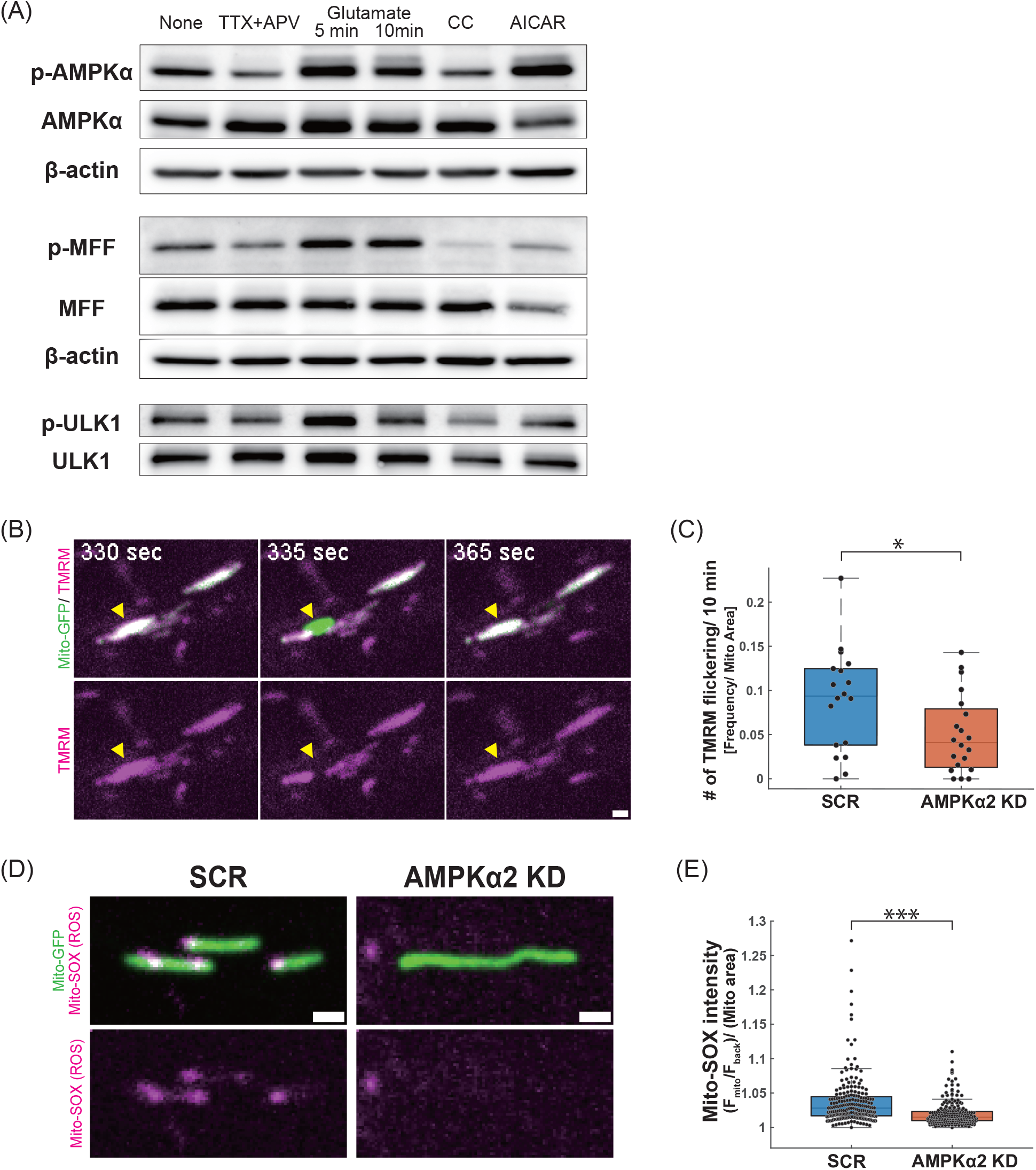
AMPK phosphorylates mitochondrial fission factor (MFF) and ULK1 in response to neuronal activity. **(A)** Phosphorylation of AMPKα and its downstream effectors in neurons treated with either TTX+APV (6h), 100 μM glutamate (5 and 10 min), 20 μM Compound C (CC, 6h) or 2 mM AICAR (6h) at DIV5. Blots were probed with antibodies against total and p-Thr172 AMPKα, total and p-Ser146 MFF, total and p-Ser555 ULK1 and β-actin. Similar results were obtained from three independent cultures. **(B)** Time-lapse images of TMRM flickering in dendritic mitochondria in hippocampal neurons. Cells were transfected with mito-GFP and SCR or AMPKα2 shRNA and treated with 20 nM TMRM. Transient loss of TMRM fluorescence in a single mitochondrion is seen (yellow arrowheads) (see also Supplementary Video 5). **(C)** Quantification of the frequencies of TMRM flickering in control and AMPKα2 KD cells. N= 18 cells for SCR, 20 cells for AMPKα2 KD. Samples were taken from four independent experiments. *p= 0.041, Wilcoxon rank-sum test. **(D)** Mito-SOX signals in dendritic mitochondria in control and AMPKα2 KD cells. Cells were transfected with mito-GFP and SCR or AMPKα2 shRNA and treated with 500 nM Mito-SOX. **(E)** Quantification of the Mito-SOX intensity in individual mitochondria. N= 212 mitochondria from 13 cells for SCR, 225 mitochondria from 17 cells for AMPKα2 KD. Samples were taken from two independent experiments. ***p= 6.9×10^−15^, Wilcoxon rank-sum test. Box plots denote the median (50th percentile) and the 25th to 75th percentiles of datasets in (C, E). Scale bars: 1 μm in (B, D).

### AMPK knockdown compromises normal mitochondrial activity in developing dendrites

We next asked if AMPK-mediated mitochondrial fission and mitophagy is required for mitochondrial quality control. Using the fluorescent dye tetramethylrhodamine methyl ester (TMRM, 20 nM), we asked whether the mitochondrial membrane potential (ΔΨm) was affected in the elongated dendritic mitochondria in neurons that were defective in AMPK-mediated mitochondrial fission. At first, we were unable to detect a significant difference in the average intensity of basal TMRM fluorescence in individual mitochondria of control and AMPKα2 knockdown cells (Extended Data Fig. 5B and C). We then analyzed dynamic changes in ΔΨm by time-lapse imaging. We observed intermittent flickers of TMRM fluorescence in some dendritic mitochondria in control neurons (Fig. 6B and Supplementary Video 5). TMRM flickering is caused by the opening of the mitochondrial permeability transition pore (mPTP), an essential process for maintaining mitochondrial homeostasis by preventing an overload of Ca^2+^ and reactive oxygen species (ROS) in the mitochondrial matrix induced by respiration activity^46,47^. In AMPKα2 knockdown neurons, the frequency of TMRM flickering was significantly reduced, suggestive of lower respiratory activity of mitochondria in these cells (Fig. 6C).

We then analyzed mitochondrial ROS levels (mitoROS) using the fluorescent dye MitoSOX (500 nM). MitoROS is a byproduct of electron transport chain activity, which is increased by high respiration and triggers mPTP opening^48^. MitoSOX signals formed puncta in the mitochondrial matrix, which were significantly decreased in AMPKα2 knockdown cells (Fig. 6D and E).

These results strongly suggest that AMPK-mediated mitochondrial fission and mitophagy is required for the maintenance of functional mitochondria in growing dendrites of hippocampal neurons.

## Discussion

To meet the rapidly growing energy demands incurred by the extension of dendritic arbors, mitochondria are actively produced and transported throughout the expanding dendritic compartments. Here we demonstrate that neuronal activity regulates mitochondrial homeostasis through the activation of the Ca^2+^-CaMKK2-AMPK axis to support dendritic outgrowth in developing neurons. AMPK deficiency prevented mitochondrial fission and mitophagy, leading to mitochondrial dysfunction with low respiratory activity and consequent growth retardation of dendrites. Our observations suggest that AMPK-dependent regulation of local mitochondrial homeostasis is an important mechanism of activity-induced dendritic growth besides the well-known transcription-dependent mechanism.

Mitochondrial fission is prerequisite for the biogenesis of mitochondria and for the elimination of damaged components through the mitophagy pathway^49,50^. Recent studies have demonstrated that symmetric fission generating two equal daughter mitochondria leads to doubling of mitochondrial biomass, while asymmetric fission generates a smaller mitochondrion that is subjected to the mitophagy pathway^45^. We observed both symmetric and asymmetric fissions in growing dendrites, but the asymmetric fission was more strongly affected by the inhibition of neuronal activity or AMPK. We showed that neuronal activity induces phosphorylation/activation of two downstream effectors of AMPK, MFF and ULK1, which are the key regulators of mitochondrial fission and mitophagy, respectively. Mitochondria with defective fission and mitophagy in AMPK-depleted neurons exhibit multiple signatures of mitochondrial dysfunction, such as reduced ΔΨm flickering and ROS production, suggesting that AMPK-mediated mitochondrial fission and mitophagy is required for the maintenance of healthy mitochondria in growing dendrites.

On the other hand, we did not obtain evidence supporting the regulation of mitochondria biogenesis by activity-induced AMPK signaling, such as a change in PGC1α phosphorylation and expression. Nonetheless, the total biomass of dendritic mitochondria should decrease as a consequence of AMPK knockdown or neuronal activity inhibition, since the mitochondrial density remains the same but the dendritic volume is substantially reduced. We will need more sensitive assays to draw conclusions regarding the regulation of mitochondrial biogenesis by activity-induced AMPK signaling.

AMPK and AMPK-related kinases are expressed in both dendrites and axons of cortical neurons, but their activities seem differentially regulated in distinct subcellular compartments. Previous studies have shown that the activities of SAD-A/B and NUAK1 and their upstream kinase LKB1 are strongly biased in the axon and regulate axon specification and branching^39,40^. The SAD kinase has been shown to regulate the fission/fusion balance of mitochondria through phosphorylation of tau in the axon^51,52^. In contrast, we demonstrate that AMPK overactivation and suppression alters mitochondrial fission, and those changes are seen only in dendrites but not in the axon (Fig. 3). The activation of AMPK in dendrites and AMPK-related kinases in the axon appears to be regulated by distinct molecular mechanisms: SAD-A/B and NUAK1 have been shown to be constitutively activated in the axon by LKB1. In contrast, AMPK phosphorylation dynamically oscillates in dendrites, depending on Ca^2+^ influx and CaMKK2 activity, while it is independent of LKB1. AMPK activity was difficult to detect in the axon due to the high basal FRET signals, presumably reflecting the high activity of SAD-A/B and/or NUAK1 in the axon. Nonetheless, neither neural activity nor AMPK overactivation enhanced fission of axonal mitochondria, suggesting that mitochondrial dynamics and transport are differentially regulated in the axon and dendrites by distinct AMPK-related kinases.

One interesting finding in the present study is that AMPK activity dynamically oscillates in synchrony with Ca^2+^ influx triggered by neuronal activity. Ca^2+^-dependent AMPK phosphorylation is rapidly downregulated in several tens of seconds, implying a regulatory loop involving a negative regulator of AMPK activity which counteracts CaMKK2-mediated phosphorylation. The molecular pathway of AMPK dephosphorylation is less understood, but a Ca^2+^-dependent phosphatase such as PP2A is a likely candidate, as PP2A has been implicated in AMPK dephosphorylation and inactivation in non-neuronal cells^53^. The dynamic oscillatory activation of AMPK would enable fine spatiotemporal tuning of mitochondrial dynamics in response to the local activity states in dendritic compartments. The controlled fission and mitophagy is critical for the maintenance of mitochondrial quality in developing dendrites, as the inhibition of AMPK-dependent mitophagy caused aberrant elongation and dysfunction of mitochondria.

Intriguingly, the CaMKK2-AMPK pathway regulating MFF and ULK2 activities has been implicated in the excessive mitochondrial fission and mitophagy in degenerating dendrites in Alzheimer’s disease models^30,31^. It is suggested that disruption of Ca^2+^ homeostasis overactivates the CaMKK2-AMPK pathway in these neurons. Thus, Ca^2+^-sensitive activation of AMPK could be a double-edged sword, promoting dendritic growth by regulating mitochondria turnover during normal development, while contributing to neurodegeneration by excessive mitochondrial degradation under pathological conditions.

It has been shown that genetic deletions of AMPK subunits exhibit little or no cell-autonomous defects in neuronal development in mice^54–57^. Conditional deletion of AMPKα1/α2 (AMPKα1–/–; AMPKα2F/F; Emx1-Cre) exhibits no overt phenotypes in the polarization and survival of cortical neurons, except for the thinning of the cortical layer at P3^29,55^. In contrast, acute treatment with AICAR or metformin induces rapid AMPK phosphorylation and disrupts axon formation in cultured cortical neurons^58^. Thus, AMPK is thought to be dispensable for neuronal differentiation during normal development, but exerts its effects when overactivated under stress conditions. In the present study, we show that defects in mitochondrial dynamics and dendrite growth by acute AMPK inhibition are evident only in differentiating neurons in the first postnatal week, and that they recovered in later stages in vivo. It is therefore possible that the transient morphological changes might have been overlooked in previous studies. Mild and transient phenotypes by chronic inhibition of AMPK might be due to the robust multi-step regulation of mitochondrial dynamics. In mature neurons, long-term potentiation (LTP) of synaptic activity causes mitochondrial fission via CaMKII and Drp1 phosphorylation^59^. The intracellular Ca^2+^ rise also induces a distinct type of mitochondrial fission initiated from the inner mitochondrial membrane^60^. AMPK might predominantly mediate spontaneous activity before synaptic maturation, while other mechanisms may take over and control mitochondrial fission in more mature neurons. Although no overt histological defects were found in mature neurons of AMPK-deficient mice, whether these neurons form and operate normal neural circuits has not been assessed. Whether AMPK has a distinct function in dendrites of mature neurons is one of many important questions that entail further study.

The present study highlights a new role of AMPK in mitochondrial homeostasis in normal development of neurons in addition to its major role in safeguarding neurons from energetic stress. Dynamic activation and inactivation of AMPK by neuronal activity are clearly important for the fine-tuning of mitochondrial homeostasis. Thus, future work should focus on the precise dynamics and mechanism of AMPK activation, as well as inactivation, for a new perspective on the physiology and pathology of AMPK signals.

## Methods

### Mice

All animals were treated in accordance with the guidelines of the Animal Experiment Committee of Kyoto University. ICR mice were kept in a 12 hr dark/light cycle at 23 ± 3°C/ 50% humidity, with standard food and water provided ad libitum, in group housing of up to three animals per cage.

### Reagents

Reagents used in this study were as follows: D, L-APV (APV, 200 μM, Sigma Aldrich, A5282), Tetrodotoxin (TTX, 0.5 μM, Abcam, ab120054), AICAR (1-2 mM, AdipoGen. Inc, AG-CR1-0061), Compound C (CC, Dorsomorphin, 20 μM, Nacalai, 18768-04), L-Glutamic acid (Glutamate, 100 μM, Sigma-Aldrich, G1251), STO-609 (10 μM, MedChem Express, HY-19805), Pim1/AKK1-IN-1 (1 μM, MedChem Express, HY-10371), TMRM (20 nM, Fujifilm, 203-18041), Mito-SOX (500 nM, Invitrogen, M36008).

### Plasmids

pCAG-EGFP, pAAV-CAG-Mito-DsRed and pAAV-CAG-Mito-EGFP were constructed as previously described^4,61^. For RNAi experiments, the target sequences for scramble control (5’-GACATTTCATCCGTTTAGTTA-3’), AMPKα1 sh#1 (5’-GGCACACCCTGGATGAATTAA-3’), AMPKα1 sh#2 (5’-GTTGTAAACCCCTATTATTTG-3’), AMPKα2 sh#1 (5’-GGTAGACAGTCGGAGCTATCT-3’), AMPKα2 sh#2 (5’-GACAATCGGAGAATAATGAAC-3’), and CaMKK2 (5’-CCCTTTCATGGATGAACGAAT-3’) were cloned into pBAsi-hH1 vector (Takara, 3220). pCAG-AMPKα1(WT) and pCAG-AMPKα2 (WT) plasmids were created by PCR amplification of cDNAs for AMPKα1 (NCBI Reference Sequence: NM_001013367.3) and AMPKα2 (NCBI Reference Sequence: NM_178143.2) from a mouse brain cDNA library and inserted into the pCAGGS vector. Resistant mutants of AMPKα1 and AMPKα2 that contained three silent mutations within the respective shRNA target sequences were generated using the Primestar mutagenesis protocol. pPBbsr2-4031NES (AMPKAR-EV) was a gift from Michiyuki Matsuda. pCAG-jRGECO1a plasmid was generated by subcloning the jRGECO1a sequence from pGP-CMV-NES-jRGECO1a (a gift from Masayuki Sakamoto) into the pCAGGS vector. The pX458-CAG-dCas9KRAB-2A-GFP plasmid was generated by subcloning the CAG promoter from pCAGGS and the dCas9KRAB sequence from CRISPR-dCas9KRAB (a gift from Kuniya Abe) into the pX458 vector (Addgene plasmid #48138). For CRISPR interference experiments, the gRNA sequence for AMPKα2 (5’-TGCCGAAGGTGCCGACGCCCAGG-3’) was inserted into pX458-CAG-dCas9KRAB-2A-GFP.

### Cell cultures and transfection

Primary cultures of hippocampal neurons were prepared as previously described with a few modifications^32^. In brief, postnatal day 0 (P0) mouse hippocampi were dissected, dissociated using Neuron Dissociation Solutions (Fujifilm, 291-78001), and plated on poly-D-lysine (Sigma-Aldrich, P6407) coated coverslips at a density of 3.0~3.5×10^5^ cells/cm^2^ in MEM (Gibco, 11095080) supplemented with 10% horse serum (Gibco, 26050070), 0.6% D-glucose, 1 mM sodium pyruvate (Sigma-Aldrich, S8636), and 1% penicillin-streptomycin. Three hours after plating, media was replaced with Neurobasal medium (Gibco, 21103049) supplemented with 2% (v/v) B-27 (Gibco, 17504044) and 0.25% (v/v) GlutaMax (Gibco, 35050061). All neurons were maintained at 37°C in 5% CO_2_. Five μM AraC (Sigma-Aldrich, C1768) was added at DIV3. Neurons were transfected at DIV3 using Lipofectamine 2000 (Thermo Fisher Scientific, 11668019) according to the manufacturer’s instructions. The CRISPRi plasmid was transfected at DIV2.

HEK293T cells (Riken BRC Cell Bank, RCB2202) were cultured in DMEM (Gibco, 11965-092) supplemented with 10% Fetal Bovine Serum (FBS, Sigma-Aldrich, F7524), 100 units/mL penicillin and 100 ug/mL streptomycin (Gibco, 15140-122) at 37°C in 5% CO_2_. Cells were passaged every 2-3 days and maintained in plastic cell culture-treated dishes. shRNA constructs were transfected using Lipofectamine 2000.

### Western blotting

Cells were harvested in RIPA lysis buffer (50 mM Tris HCl (pH 7.4), 150 mM sodium chloride, 0.25% (w/v) Sodium Deoxycholate, 1% (v/v) NonidetP-40, 1 mM EDTA, and Protease and phosphatase inhibitor cocktail (Thermo Fisher Scientific, 78442)). Lysates were subjected to SDS-PAGE and immunoblotting.

Primary antibodies used were the following: mouse anti-GFP (1:5000, Thermo Fisher Scientific, A11120); rabbit anti-AMPKα1 (1:3000, Abcam, ab32047); rabbit anti-AMPKα2 (1:3000, Abcam, ab3760); rabbit anti-AMPKα (1:1000, Cell Signaling Technology, 2532); rabbit anti-phospho-AMPKα (Thr172) (1:1000, Cell Signaling Technology, 2535); rabbit anti-MFF (1:3000, Proteintech, 17090-1-AP); rabbit anti-phospho-MFF (Ser172/146) (1:1000, Affinity Biosciences, AF2365); rabbit anti-ULK1 (1:2000, Cell Signaling Technology, 8054); rabbit anti-phospho-ULK1 (Ser555) (1:1000, Cell Signaling Technology, 5869); rabbit anti-PGC1α (1:5000, Abcam, ab191838) and HRP-conjugated mouse anti-beta actin (1:10000, Santa Cruz, sc-47778 HRP). The secondary antibodies used were the following: HRP-conjugated goat anti-mouse IgG (1:10000, Bio-Rad, 170-6516); HRP-conjugated goat anti-rabbit IgG (1:10000, Bio-Rad, 170-6515).

### In utero electroporation

Pregnant mice on day 14.5 of gestation were deeply anesthetized via the intra-abdominal injection of a mixture of medetomidine, midazolam and butorphanol. Plasmid DNA diluted with saline containing 0.01% Fast Green was injected into the lateral ventricle of mouse embryos with a glass needle. Four current pulses (amplitude, 40 V; duration, 50 ms; intervals, 950 ms) were delivered with a forceps-shaped electrode (CUY650P3, NepaGene) connected to an electroporator (CUY21, NepaGene). Concentrations of the plasmids used are as follows: pCAG-EGFP (0.2 μg/μl); shRNA-scramble control (3-6 μg/μl); shRNA-AMPKα2 (3-6 μg/μl); CRISPRi gRNA-empty (5 μg/μl); and CRISPRi gRNA-AMPKα2 (5 μg/μl).

### Immunofluorescence and image acquisition

For immunocytochemistry, cultured cells were fixed with 4% PFA in PBS and permeabilized with PBS containing 0.25% Triton X-100 (PBS-T). Cells were then blocked with blocking solution (PBS with 2% BSA) and incubated with rabbit anti-AMPKα2 primary antibody (1:200, Proteintech, 18167-1-AP) at 4°C overnight. After washing with PBS, cells were incubated with goat anti-Rabbit Alexa Fluor 647 (1:400, Invitrogen, A21244) secondary antibody at 4°C overnight.

For immunohistochemistry, the mice at P5 or P10 were perfused with PBS followed by 4% paraformaldehyde (PFA) in phosphate buffer. Their brains were removed and postfixed for 2-3 hrs at 4°C. After washing with PBS, the brains were embedded in 3.5% low-melting agarose in PBS. The brains were then sectioned into 200 μm thick sagittal slices with a vibratome (NLS-AT, Dosaka EM). The sections were permeabilized in PBS with 0.25 % Triton (PBS-T) for 15 min and blocked with 2% skim milk in PBS-T for 30 min. The sections were incubated with a chick anti-GFP (1:2000, Invitrogen, A10262) primary antibody in blocking solution at 4°C overnight, followed by incubation with a goat anti-Chicken Alexa Fluor 488 (1:400, Invitrogen, A11039) secondary antibodies at 4°C overnight.

Images of the fixed samples were acquired with a laser scanning confocal microscope (FV1000, Olympus) equipped with UPlanSApo 20× (N.A. 0.75), UPLSAPO 40× (NA 0.95) objectives, or a spinning disk confocal microscope (Dragonfly, Andor) equipped with Nikon PLAN APO 40× (Silicone oil, NA 1.25), Apo TIRF 100× (oil-immersion, NA 1.49) objectives.

For morphometric analysis, dendrites were traced using the ImageJ plugin Simple Neurite Tracer (SNT). Total dendrite length and the number of branches were calculated using MATLAB software. For quantification of dendritic mitochondrial densities (ratio of mitochondrial area to dendrite area), acquired images were binarized and the areas filled with Mito-DsRed signals and volume marker GFP signals were quantified using ImageJ.

For p62 analysis, hippocampal neurons transfected with GFP and mito-DsRed were fixed at DIV5 and immunostained with guinea pig anti-p62 primary antibody (1:100, Progen, GP62-C,) and goat anti-Guinea Pig Alexa Fluor 633 secondary antibody (1:400, Invitrogen, A21105). The number of p62 puncta in dendrites was counted using the ImageJ plugin Find Maxima. For colocalization analysis of p62 puncta and mitochondria, Mander’s overlap coefficient was calculated from anti-p62 signals and mito-DsRed signals in dendrites using the ImageJ plugin JaCoP. Surface rendering was done with Imaris software.

### Time-lapse imaging and image analysis

For time-lapse imaging of mitochondrial dynamics, cultured hippocampal neurons were transfected on DIV3 with pCAG-EGFP and pAAV-CAG-mitoDsRed plasmids. Time-lapse images were acquired every 3 s for 20 min at DIV5 using a spinning-disc confocal microscope (CV1000, Yokogawa) with a UPLSApo 100× objective (Oil immersion, N.A. 1.4, Olympus) at 37°C with 5% CO_2_ flow. The mitochondrial movement was tracked using the ImageJ plugin MTrackJ. The mitochondrial fission and fusion events were counted manually. Peripheral and midzone mitochondrial fissions were defined as events in which the relative position of the fission site along the length axis was 0~25% and 25~50%, respectively. The percentage of motile mitochondria was calculated by dividing the number of mitochondria that were displaced more than 4 μm from their original positions within 20 minutes by the total number of mitochondria. A continuous movement of mitochondria (> 1.5 μm for > 6 sec) was defined as one moving event, and their speed was calculated by using MATLAB software.

For AMPK FRET imaging, cultured hippocampal neurons transfected with AMPKAR-EV were imaged every 3 sec for 3min at DIV5 with an inverted epifluorescence microscope (IX83, Olympus) equipped with LUCPlanFLN 20× (NA 0.45), UplanSApo 40× (Silicone oil, NA 1.25) objectives. For dual imaging of AMPK FRET and calcium imaging, cultured hippocampal neurons were cotransfected with AMPKAR-EV and jRGECO1a, and imaged every 5 sec for 5 min. Fluorescence images were acquired with the following filters and mirrors: excitation filters FF02-438/24-25 (for CFP/ YFP); a dichroic mirror FF458-Di02-25×36 (for CFP/ YFP); emission filters FF01-483/32-25 (for CFP); FF01-542/27-25 (for YFP); and filter sets U-FMCHE (Ex 565-585 nm/ DM 595/ Em 600-690) (for jRGECO1a).

For FRET ratio analysis, after background subtraction, the YFP/CFP fluorescence intensity ratio was calculated in the ROI of the proximal dendritic region. The YFP/CFP ratio was plotted along the time axis, and the FRET peak frequency was quantified using the MATLAB findpeaks function with a threshold for peak heights greater than 0.05. The Cross-Correlation Function was calculated followed methods described previously ^62^ using MATLAB software.

For TMRM imaging, cultured hippocampal neurons transfected with mito-EGFP were incubated with 20 nM TMRM 1-2 h prior to imaging. Time-lapse images were acquired every 5 sec for 10 min at DIV5 with a spinning-disc confocal microscope (CV1000, Yokogawa) equipped with a UPLSApo 100× objective (Oil immersion, N.A. 1.4, Olympus).

For TMRM intensity analysis, after background subtraction, TMRM intensity in individual mitochondria was divided by the mitochondrial area. TMRM flickering, defined as a transient loss of TMRM intensity to the background level, was scored as the number of flickering events that occurred within 10 minutes in the soma and proximal dendrites, divided by the mitochondrial area.

For quantification of mitochondrial ROS, cultured hippocampal neurons transfected with mito-EGFP at DIV3 were incubated with 500 nM Mito-SOX for 15 min at DIV5 and then the Mito-Sox was washed out. Time-lapse imaging was initiated 18-20 h after treatment, using a spinning-disc confocal microscope (CV1000, Yokogawa) equipped with a UPLSApo 100× objective (Oil immersion, N.A. 1.4, Olympus). Mito-SOX intensity in individual mitochondria ((F_mitochondria_/ F_background_)/ (Mitochondrial Area)) was quantified using ImageJ.

## Acknowledgements

We thank Dr. Michiyuki Matsuda for the AMPKAR-EV plasmid, Dr. Yoshiaki Tagawa for the GCaMP6s plasmid, Dr. Masayuki Sakamoto for the jRGECO1a plasmid, Drs. Kuniya Abe, Yusuke Kishi, Shinnosuke Suzuki, Ryuta Kinoshita for the CRISPR-dCas9KRAB plasmid, iCeMS Analysis Center for technical assistance with microscopy, Drs. Takaki Miyata, Mayumi Okamoto, Daniel Packwood for advice, Dr. James Hejna for critical reading of the manuscript.

## Author contributions

Conceptualization: A.H., K.F., M.K.; Methodology: A.H., K.F., A.K., N.O., M.K.; Investigation: A.H., J.K., K.F.; Writing - original draft: M.K. and A.H., and all the authors read and commented on the manuscript.

---

**Extended Data Figure 1.**
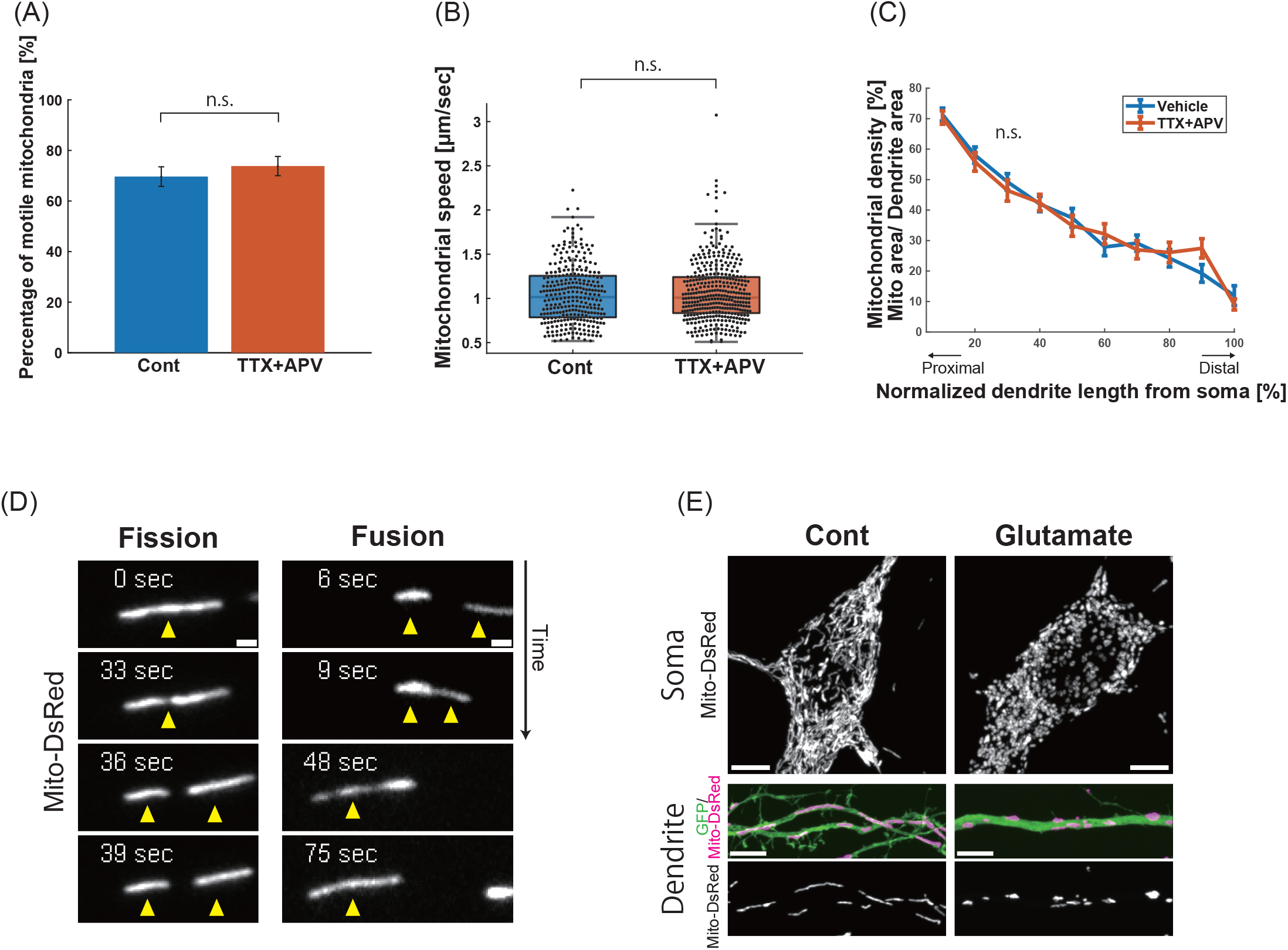
Neuronal activity regulates mitochondrial dynamics. **(A)** Motile fraction of mitochondria in dendrites before and after TTX+APV treatment. Cells were labeled with mito-DsRed and GFP. N= 14 cells, mean ± SEM, (n.s.) p= 0.45, unpaired two-tailed t test. **(B)** Mitochondrial speed before and after TTX+APV treatment. N= 294 events from 14 control cells, 376 events from 14 drug-treated cells, (n.s.) p= 0.52, Wilcoxon rank-sum test. **(C)** Mitochondrial distribution along the shaft of primary dendrites. The primary dendrites were compartmentalized by the relative length from the soma, with the tip of the dendrite as 100%. Hippocampal neurons expressing GFP and mito-DsRed were treated with vehicle or TTX+APV for 2 days from DIV3 and fixed at DIV5. Mitochondrial density was calculated by dividing the mitochondrial area (demarcated by mitoDsRed) by the dendritic area (demarcated by GFP). N= 23 control cells, 30 drug-treated cells, (n.s.) p> 0.05, unpaired two-tailed t test. **(D)** Time-lapse images of fission and fusion of dendritic mitochondria in hippocampal neurons labeled with GFP and mito-DsRed (see also Supplementary Video 3). **(E)** Mitochondrial morphology in soma and dendrites after 1-min treatment with vehicle or 100 μM Glutamate at DIV6. Scale bars: 1 μm in (D); 5 μm in (E). Samples were taken from three independent experiments.

**Extended Data Figure 2.**
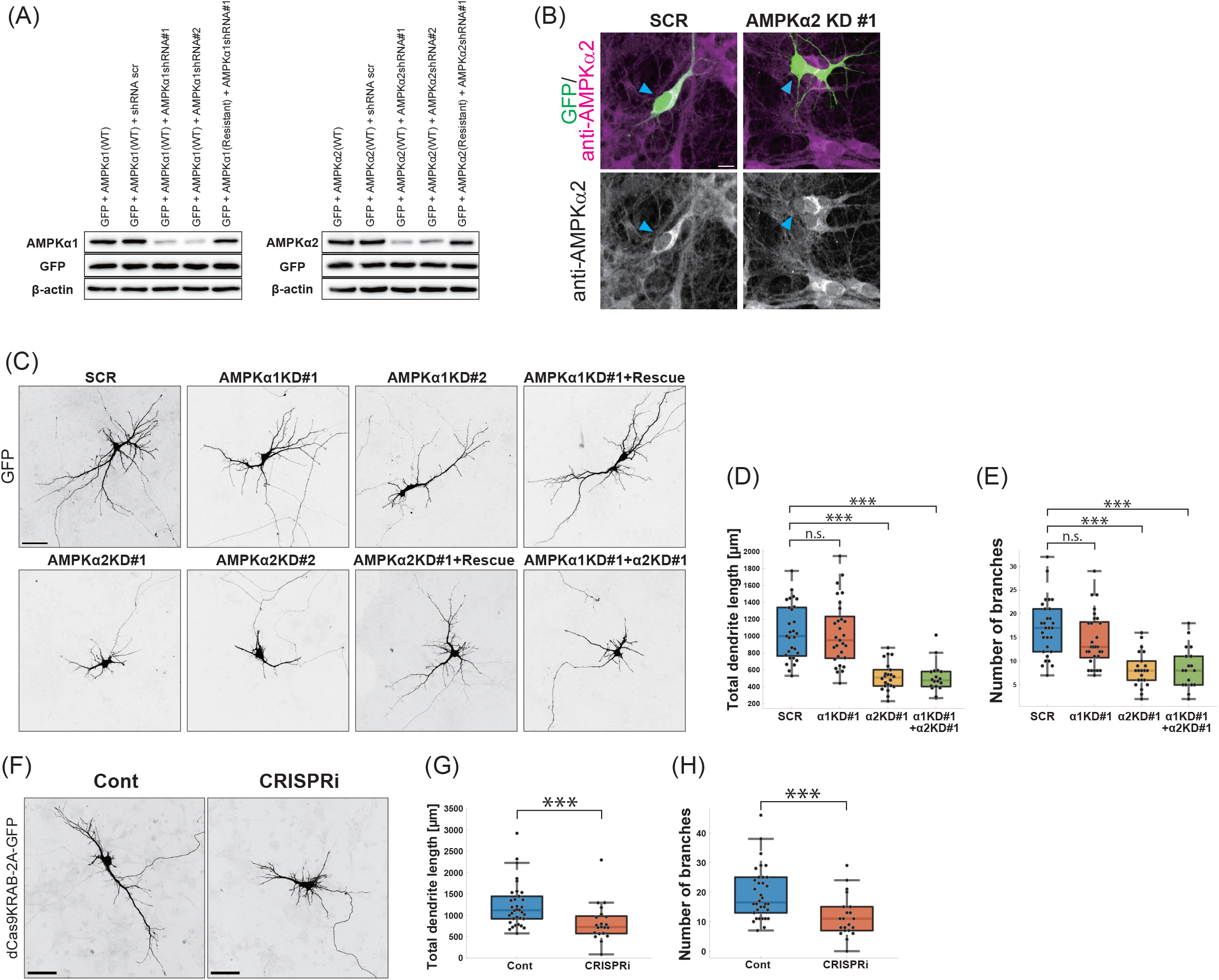
Validation of shRNA targeting AMPKα subunits. **(A)** HEK293T cells were transfected with GFP and AMPKα 1 or AMPKα2 cDNA plasmids with or without respective shRNA constructs. Cell lysates were prepared 2 days after transfection and analyzed by western blotting with antibodies against GFP, AMPKα1, AMPKα2 and β-actin. **(B)** DIV5 hippocampal neurons transfected with GFP and shRNA for the scrambled control (SCR) or AMPKα2 (α2 KD #1) were immunostained with an antibody against AMPKα2. Arrowheads indicate shRNA-transfected neurons. **(C)** Representative images of DIV5 hippocampal neurons transfected with GFP and shRNA for the scrambled control (SCR), AMPKα1 (α1 KD #1 or #2), AMPKα2 (α2 KD #1 or #2), AMPKα1 and α2 (α1KD #1 + α2KD #1), or AMPKα1 (α1 KD #1) or AMPKα2 (α2 KD #1) together with the respective shRNA-resistant mutants (Rescue). **(D, E)** Quantification of total dendritic length (D) and number of branch points (E) of hippocampal neurons described in (C). N= 29 cells for SCR, 29 cells for α1 KD #1, 22 cells for α2 KD #1 from three independent experiments, and 18 cells for α1KD #1 + α2KD #1 from two independent experiments; ***p<0.001; (n.s.) p> 0.05; Kruskal–Wallis test with Bonferroni multiple comparisons test. **(F)** Morphology of hippocampal neurons transfected with CRISPR interference plasmids for gRNA-empty control (Cont) or gRNA-AMPKα2 (AMPKα2 CRISPRi). Cells were transfected at DIV2 and fixed at DIV5. **(G, H)** Quantification of total dendritic length (G) and number of branch points (H) of neurons described in (F). N= 34 cells for control, 21 cells for AMPKα2 CRISPRi from three independent experiments. ***p= 9.7×10^−4^ (G), ***p= 6.4×10^−4^ (H), Wilcoxon rank-sum test. Box plots denote the median (50th percentile) and the 25th to 75th percentiles of datasets in (D, E, G, H). Scale bars: 10μm in (B), 50 μm in (C, F).

**Extended Data Figure 3.**
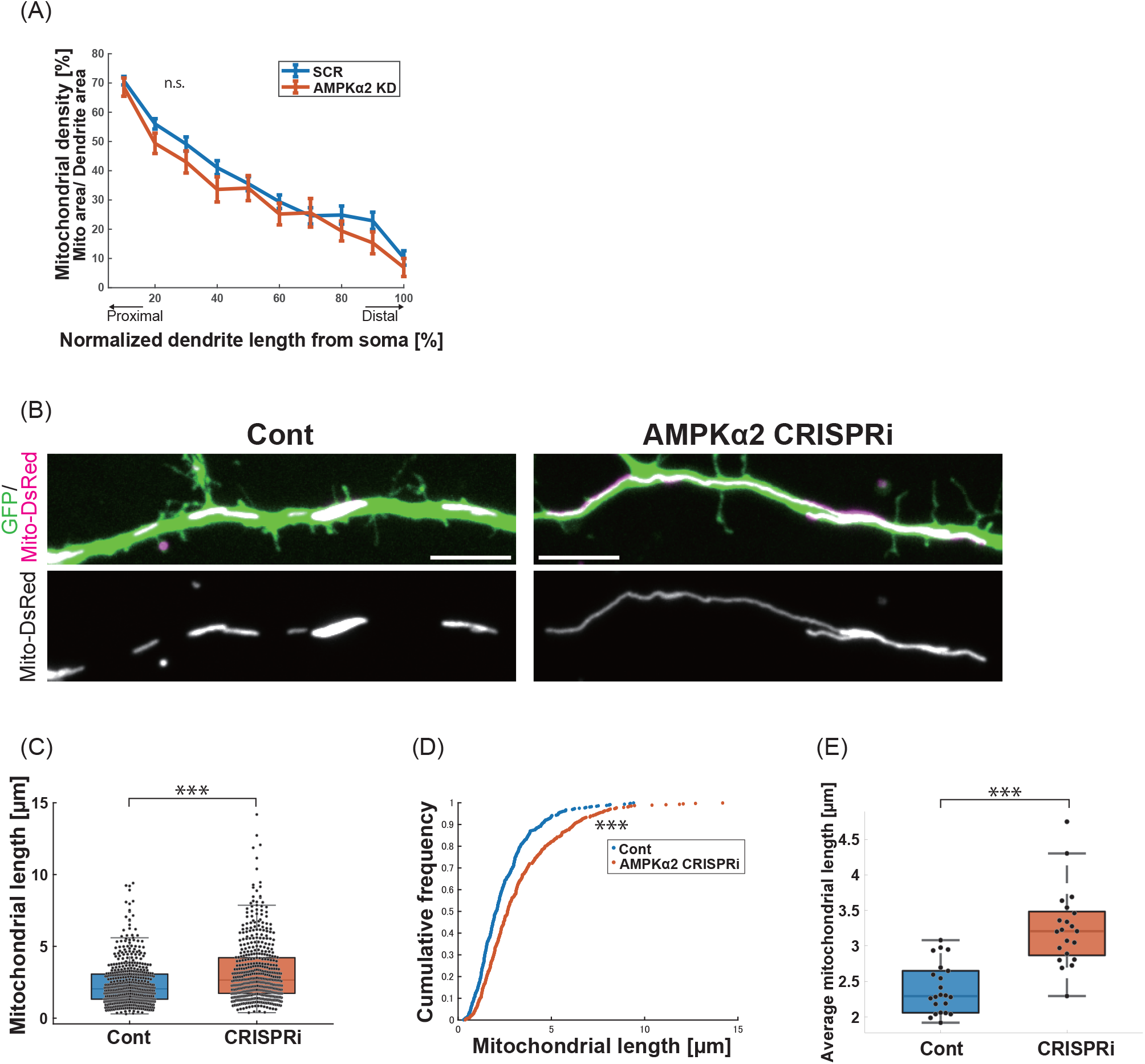
AMPK induces mitochondrial fission in dendrites of growing hippocampal neurons. **(A)** Mitochondrial distribution along the shaft of primary dendrites. Hippocampal neurons were transfected with GFP, mito-DsRed and shRNA-scramble control (SCR) or shRNA-AMPKα2 (AMPKα2 KD) at DIV3 and fixed at DIV5. The primary dendrites were compartmentalized by the relative length from the soma, with the tip of dendrite defined as 100%. The mitochondrial density was calculated by dividing the mitochondrial area (demarcated by mitoDsRed) by the dendritic area (demarcated by GFP). N= 22 cells for SCR, 24 cells for AMPKα2 KD; (n.s.) p> 0.05; unpaired two-tailed t test. **(B)** Morphology of dendritic mitochondria in hippocampal neurons transfected at DIV2 with CRISPR interference plasmids for gRNA-empty control (Cont) or gRNA-AMPKα2 (AMPKα2 CRISPRi) and fixed at DIV5. Cells were labeled with mito-DsRed and GFP. Scale bars: 5 μm. **(C, D)** Distribution (C) and cumulative frequency (D) of the mitochondrial length in dendrites. N= 534 mitochondria from 22 control cells, 497 mitochondria from 21 CRISPRi cells; *** p= 8.0×10^−12^; Wilcoxon rank-sum test. **(E)** Average mitochondrial length in individual cells. N=22 cells for control, 21 cells for CRISPRi; *** p= 2.3×10^−6^; Wilcoxon rank-sum test. Box plots denote the median (50th percentile) and the 25th to 75th percentiles of datasets in (C, E). Samples were taken from three independent experiments.

**Extended Data Figure 4.**
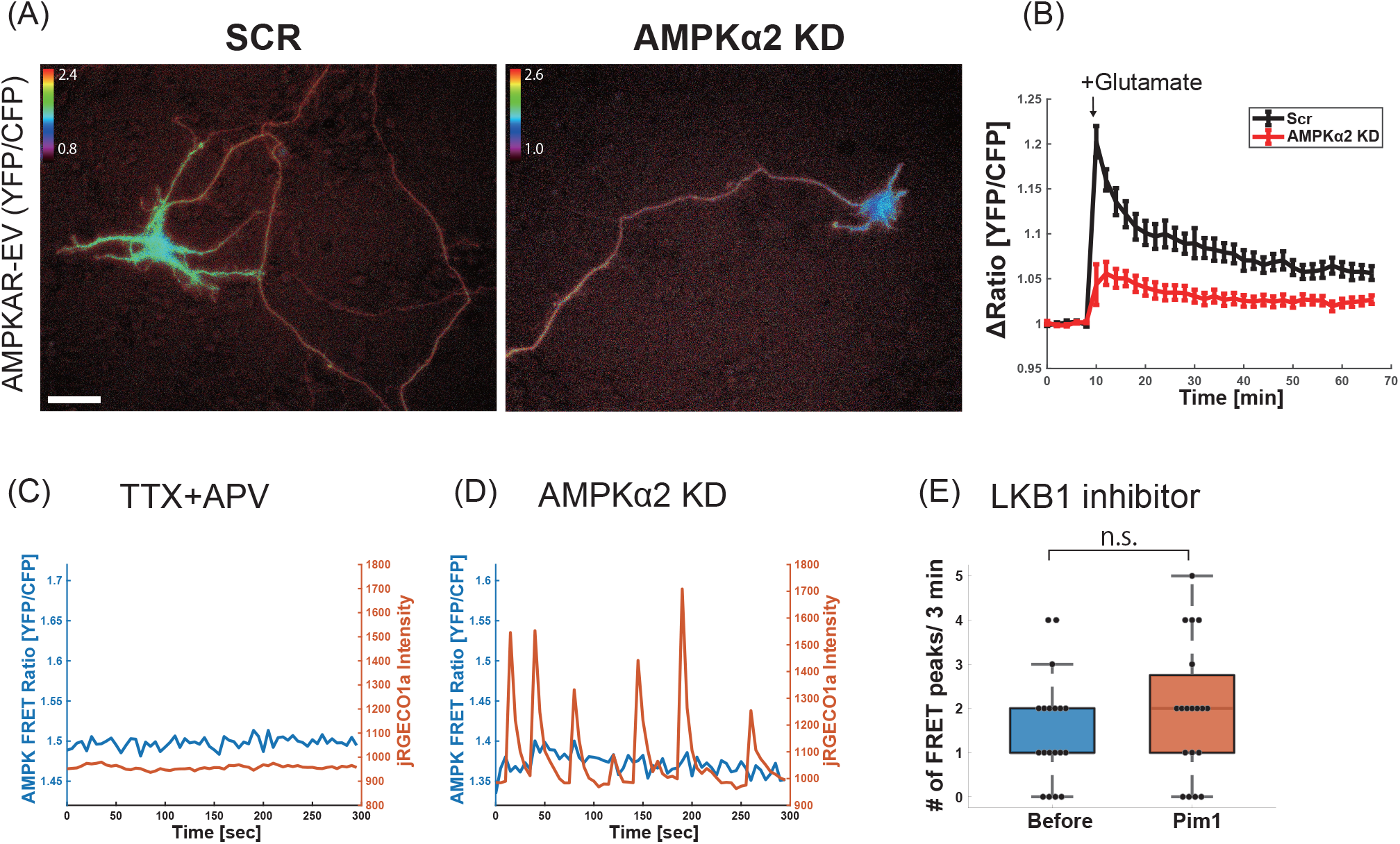
AMPK activity in hippocampal neurons visualized by AMPKAR-EV. **(A)** Subcellular distribution of AMPKAR-EV FRET signals in hippocampal neurons at DIV5. Cells were transfected with AMPKAR-EV together with shRNA-scramble control (SCR) or shRNA-AMPKα2 (AMPKα2 KD). Scale bar: 50 μm. **(B)** Activation of AMPK induced by neuronal activity. DIV5 hippocampal neurons expressing AMPKAR-EV with shRNA-scramble control (SCR) or shRNA-AMPKα2 #1 (AMPKα2 KD) were treated with 1 μM glutamate. The YFP/CFP ratio in dendrites was monitored every 2 min for 70 min before and after glutamate treatment. N= 18 cells for control and AMPKα2 from two independent experiments. **(C)** Representative traces of AMPK activity and Ca^2+^ influx in hippocampal neurons treated with TTX+APV. Cells were transfected with AMPKAR-EV and jRGECO1a. **(D)** Representative traces of AMPK activity and Ca^2+^ influx in hippocampal neurons transfected with AMPKAR-EV, jRGECO1a and shRNA-AMPKα2. **(E)** Quantification of the frequency of AMPKAR-EV FRET peaks in hippocampal neurons expressing AMPKAR-EV before and after treatment with 1 μM Pim1 (LKB1 inhibitor) at DIV5. N= 19 cells from two independent experiments, (n.s.) p= 0.30, Wilcoxon signed-rank test. Box plots denote the median (50th percentile) and the 25th to 75th percentiles of datasets.

**Extended Data Figure 5.**
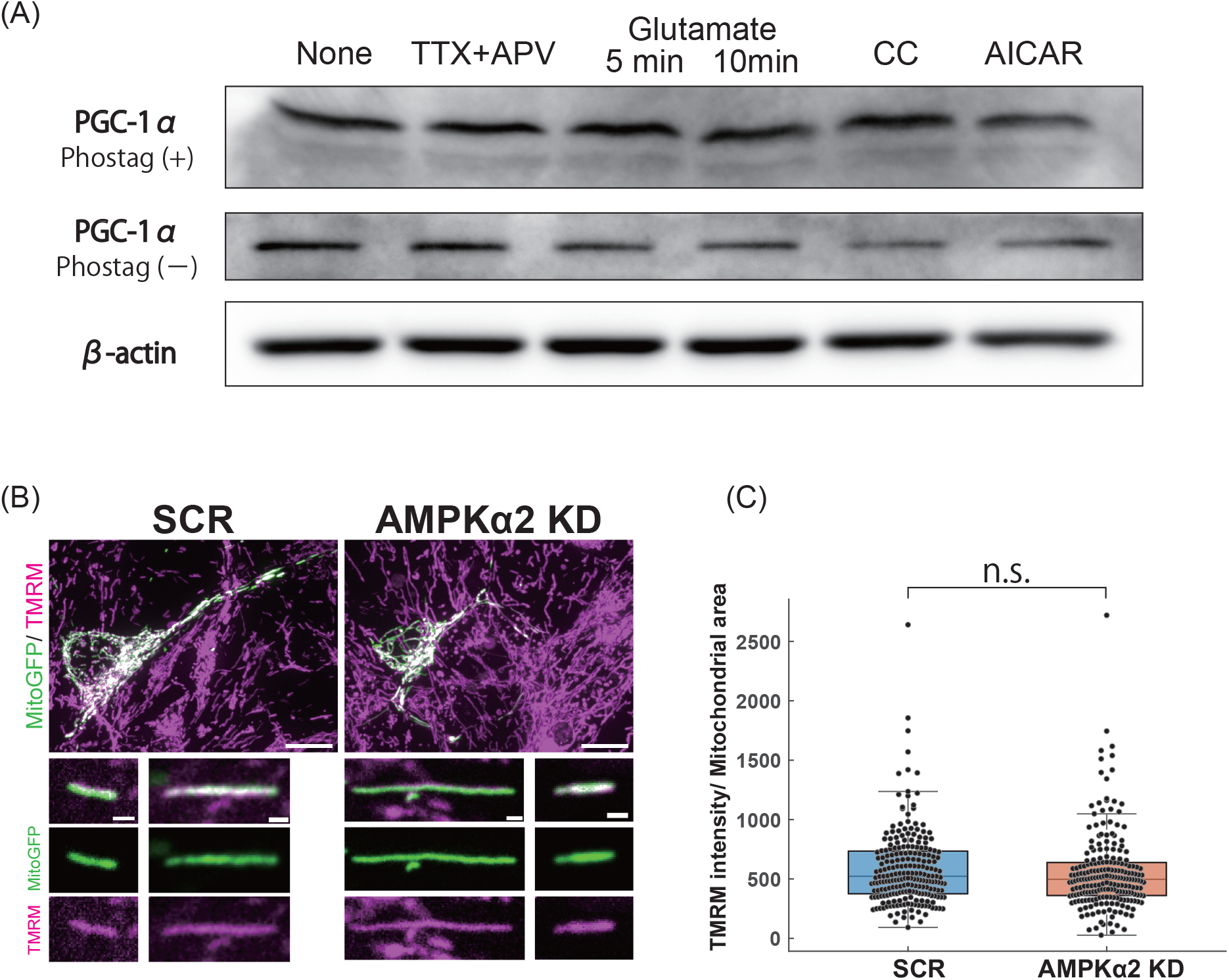
Effects of AMPK deficiency on PGC1α phosphorylation and mitochondrial membrane potential. **(A)** Phos-tag SDS-PAGE for a mitochondrial biogenesis marker, PGC1α in neurons treated with either TTX+APV (6h), 100 μM glutamate (5 or 10 min), 20 μM Compound C (CC, 6h), or 2 mM AICAR (6h). Cell lysates were subjected to Phos-tag SDS-PAGE (100 μM Phos-tag) and standard SDS-PAGE, followed by immunoblotting with antibodies for PGC1α and β-actin. **(B)** Representative images of dendritic mitochondria in neurons treated with 20 nM TMRM (magenta). Cells were transfected with mito-GFP (green) together with shRNA-scramble control (SCR) or shRNA-AMPKα2 (AMPKα2 KD). **(C)** Quantification of TMRM intensity in individual mitochondria. N= 227 mitochondria from 17 control cells, 216 mitochondria from 15 AMPKα2 KD cells from two independent experiments. (n.s.) p= 0.14, Wilcoxon rank-sum test. Box plots denote the median (50th percentile) and the 25th to 75th percentiles of datasets. Scale bars: 10 μm (B, upper panels), 1 μm (B, lower panels).

## Supplementary information

**Supplementary Video 1. Calcium transients in hippocampal neurons before and after TTX+APV treatment.** Cells were transfected with GCAMP6s at DIV3. Images were acquired every 3 sec for 5 min at DIV5 using spinning disk confocal microscopy. Display rate: 10 frames/second. Scale bar: 10 μm. Related to Fig. 1A.

**Supplementary Video2. Time-lapse movies of growing hippocampal neurons cultured in the presence or absence of TTX+APV.** Cells were labeled with GFP. Images were acquired every 30 min for 52 h from DIV5 to 7 using spinning disk confocal microscopy. Display rate: 10 frames/second. Scale bar: 50 μm. Related to Fig. 1B.

**Supplementary Video3. Time-lapse movies of mitochondrial fission and fusion in dendrites** Cells were labeled with mito-DsRed and GFP. Time-lapse images were acquired every 3 sec for 20 min at DIV5 using spinning disk confocal microscopy. Display rate: 8 frames/second. Scale bar: 1 μm. Related to Extended Data Fig. 1D.

**Supplementary Video 4. Simultaneous imaging of AMPK activity and Ca^2+^ transients before and after TTX+APV treatment.**

Cultured hippocampal neurons were transfected with AMPKAR-EV and jRGECO1a. The pseudo-color represents the YFP/CFP ratio (AMPK activity, left), and the Ca^2+^ influx (right), respectively. Images were acquired every 5 sec for 5 min at DIV5 using epifluorescence microscopy. Display rate: 5 frames/second. Scale bar: 10 μm. Related to Fig. 4E-G.

**Supplementary Video 5. Time-lapse movie of mitochondrial TMRM flickering.**

Cultured hippocampal neurons transfected with mito-GFP (green) were treated with 20 nM TMRM (magenta) at DIV5. Images were acquired every 5 sec for 10 min using spinning disk confocal microscopy. Display rate: 8 frames/ second. Scale bar: 1 μm. Related to Fig. 6B.

